# Mitochondrial structural and functional defects in the *Drosophila melanogaster* model of PLA2G6 Associated Neurodegeneration (PLAN)

**DOI:** 10.64898/2026.02.21.707236

**Authors:** Rubaia Tasmin, Devisri Pranvi Matam, Surya Jyoti Banerjee

## Abstract

PLA2G6-associated neurodegeneration (PLAN) is a rare progressive disorder caused by mutations in *PLA2G6*, which encodes calcium-independent phospholipase A2, an enzyme required for phospholipid remodeling and membrane lipid homeostasis through the Lands cycle. Although mitochondrial dysfunction has been implicated in PLAN, how *PLA2G6* loss affects mitochondrial structure and function across tissues, age, and sex remains unclear. Here, we used homozygous null iPLA_2_-VIA mutant *Drosophila melanogaster*, a model of PLAN, to examine mitochondrial ultrastructure, abundance, function, DNA content, and mitochondrial maintenance gene expression. Transmission electron microscopy revealed severe mitochondrial abnormalities in the brain, thorax, and ovary from 7-day-old and 3-week-old mutant flies, including disrupted cristae, abnormal morphology, and compromised membrane structure. Mutants also showed reduced mitochondrial number, which became more widespread with age. MitoTracker imaging further showed reduced mitochondrial staining and abnormal clumped mitochondrial distribution in mutant brains. Consistent with mitochondrial depletion, mitochondrial DNA content was reduced in mutants, with decreased *ATPase6* but unchanged nuclear *ATPSynC* levels. Functionally, mutants exhibited reduced ATP production across multiple tissues and at the whole-body level, while ROS levels changed in a tissue-, age-, and sex-dependent manner. Transcript analysis showed reduced *mTOR* and *PGC-1α* expression, altered expression of fusion and fission genes including *Opa1*, *Drp1*, and *Fis1*, age- and sex-dependent changes in *Pink1*, reduced *Trap1* expression in aged mutants, and unchanged *Sirtuin 6* expression. Together, these findings show that iPLA_2_-VIA is required for mitochondrial maintenance and bioenergetic integrity in vivo.

## Introduction

Mitochondria are essential subcellular organelles found in eukaryotic cells, where they support cellular metabolism and ATP (Adenosine Triphosphate) production (Beck et al. 2011). They are defined by an outer and an inner membrane, the intermembrane space, and the mitochondrial matrix (McBride et al. 2006). The outer membrane contains pore-forming channels that allow the movement of selected metabolites and ions between the cytoplasm and mitochondria (Nelson and Kabir 1986). The inner membrane is extensively folded into cristae within the matrix (McBride et al. 2006). These cristae provide the structural platform for the electron transport chain and ATP synthesis by oxidative phosphorylation (Beck et al. 2011). During oxidative phosphorylation, electrons released from NADH (reduced Nicotinamide Adenine Dinucleotide) and FADH_2_ (reduced Flavin Adenine Dinucleotide) enter the electron transport chain through Complexes I and II and are then transferred through Complex III and Complex IV. Complexes I, III, and IV pump protons across the inner mitochondrial membrane into the intermembrane space. This creates an electrochemical gradient that ATP synthase (Complex V) uses to generate ATP (Quintana-Cabrera et al. 2018). However, when electron transfer is disrupted, electrons can leak from the electron transport chain, particularly from Complexes I and III, and react with oxygen to form reactive oxygen species (ROS). At low levels, ROS participates in cellular signaling. However, excessive or poorly controlled ROS can damage mitochondrial lipids, proteins, and DNA, further impairing mitochondrial function and promoting neurodegeneration (Zorov et al. 2014). The mitochondrial *Superoxide dismutase 2* (*Sod2*) is a mitochondrial matrix antioxidant enzyme that protects mitochondria from oxidative damage by converting superoxide radicals (O₂•⁻) into hydrogen peroxide (H₂O₂) (Islam et al. 2026). In addition to supporting oxidative phosphorylation, the inner mitochondrial membrane is enriched in cardiolipin, a phospholipid that helps maintain cristae architecture and supports the stability and function of respiratory chain complexes (McBride et al. 2006). Therefore, intact mitochondrial membranes, appropriate membrane potential, and well-preserved cristae are essential for efficient ATP production and neuronal energy homeostasis (Quintana-Cabrera et al. 2018).

Mitochondrial function also depends on the coordinated regulation of mitochondrial production, maintenance, and turnover. One major pathway involved in mitochondrial production is mitochondrial biogenesis. The mechanistic Target of Rapamycin (mTOR) promotes mitochondrial biogenesis by increasing the expression of *spargel* (*srl*), the *Drosophila* ortholog of human *PGC-1α* (*peroxisome proliferator-activated receptor gamma coactivator-1 alpha*) (Morita et al. 2013). PGC-1α is considered the central transcriptional co-activator of mitochondrial biogenesis (Wenz 2011). When mTOR activity is reduced, *PGC-1α* levels and activity often decrease. This can lead to reduced expression of mitochondrial genes and reduced mitochondrial DNA replication. Therefore, reduced *mTOR–PGC-1α* signaling can limit mitochondrial biogenesis and contribute to decreased mitochondrial abundance and function (Cunningham et al. 2007).

Along with biogenesis, mitochondrial dynamics are essential for maintaining mitochondrial structure and function. Mitochondrial fusion is a two-step process that first joins the outer membranes and then the inner membranes of two mitochondria. This process allows mitochondria to mix their contents, maintain mitochondrial DNA integrity, and support efficient energy production. Mitochondrial fusion is regulated by the coordinated actions of fuzzy onions (fzo) and Mitochondrial assembly regulatory factor (Marf), the *Drosophila* orthologs of human Mitofusion 1 and 2 (Mfn1 *and* Mfn2), respectively, at the outer mitochondrial membrane. Fusion also requires optic atrophy 1 (Opa1) at the inner mitochondrial membrane. These GTPase proteins mediate the tethering and merging of neighboring mitochondria, promoting the exchange of mitochondrial contents and preserving mitochondrial network integrity (Jia et al. 2025). Disruption of any of these proteins can shift mitochondria toward fragmentation and structural dysfunction (Chen et al. 2005). Mitochondrial fission is the process by which a single mitochondrion divides into two or more daughter organelles. Dynamin-related protein 1 (Drp1) is the main fission protein. It is a cytosolic GTPase that is recruited to the outer mitochondrial membrane at specific constriction sites. Once recruited, Drp1 assembles into ring-like spirals around the mitochondrion. Using energy from GTP hydrolysis, these spirals tighten and physically split the organelle. Fission, mitochondrial 1 (Fis1) is an outer mitochondrial membrane protein that helps recruit and anchor Drp1 to mitochondria. It acts as an adaptor or docking factor that promotes Drp1 localization to fission sites (Seo et al. 2020). Proper Drp1–Fis1 activity ensures balanced mitochondrial division, quality control, and removal of damaged mitochondria.

In addition to biogenesis and dynamics, mitochondrial quality control also depends on mitophagy, the selective removal of damaged mitochondria. PTEN-induced putative kinase 1 (Pink1) is a key regulator of this pathway. Pink1 is a mitochondrially targeted serine/threonine kinase linked to mitochondrial quality control and neurodegeneration. When mitochondria lose membrane potential, Pink1 accumulates on the outer mitochondrial membrane and helps recruit Parkin, marking damaged mitochondria for removal through mitophagy (Chu 2010; de Vries and Przedborski 2013; Pickrell and Youle 2015). Mutations in human Pink1 and Parkin cause autosomal recessive early-onset Parkinson’s disease. These mutations disrupt mitochondrial quality control by impairing the recognition and mitophagic removal of damaged mitochondria. This can promote the accumulation of dysfunctional mitochondria, reduce ATP production, induce oxidative stress, and increase vulnerability of dopaminergic neurons to degeneration (Borsche et al. 2021; Corti et al. 2011; McInerney-Leo 2005; Valente et al. 2004). Tumor necrosis factor receptor-associated protein 1 (Trap1) is another mitochondrial quality-control protein linked to mitochondrial stress protection and neurodegenerative disease. In *Drosophila*, Trap1 protects against mitochondrial dysfunction in Pink1/Parkin Parkinson’s disease models. Neuronal *Trap1* upregulation rescues mitochondrial impairment in *Pink1* mutants, whereas loss of *Trap1* reduces mitochondrial function and ATP levels (Costa et al. 2013). *Sirtuin 6 (Sirt6)* is a mitochondrial stress-related regulator that supports brain mitochondrial function and metabolic homeostasis. Sirt6 deficiency has been linked to reduced mitochondrial gene expression, altered TCA cycle metabolites, decreased mitochondrial number, impaired membrane potential, and increased ROS production (Guo et al. 2022; Smirnov et al. 2023). These changes are especially relevant to neurodegeneration because similar mitochondrial and Sirt6-related alterations have been reported in aging brains and in Alzheimer’s, Parkinson’s, Huntington’s, and amyotrophic lateral sclerosis disease (Smirnov et al. 2023).

Neurodegenerative diseases (NDs) represent a major and growing global health concern and are characterized by the progressive and irreversible loss of neuronal structure and function (Gadhave et al. 2024). Common NDs such as Alzheimer’s, Parkinson’s, and Huntington’s diseases differ in their clinical manifestations, yet they share several underlying cellular mechanisms that lead to neuronal damage (Yang 2025), including oxidative stress, mitochondrial dysfunction and damage, which together drive a self-propagating cascade of neuronal damage and death (Jurcau 2021; Liu et al. 2024). Neurons have exceptionally high energy demands, and ATP generated by mitochondrial oxidative phosphorylation is essential for neuronal maintenance, synaptic activity, and regeneration (Ren et al. 2024). Mitochondrial function depends mostly on intact membrane architecture and well-organized cristae, which maximize the surface area available for the electron transport chain (Quintana-Cabrera et al. 2018). Consequently, alterations in mitochondrial structure, abundance, and cellular distribution can directly compromise neuronal viability. These mitochondrial defects are increasingly recognized as central contributors to age-related and genetic mutation-driven neurodegeneration, like in PLA2G6-associated neurodegeneration (PLAN) (Banerjee et al. 2021b; Beck et al. 2011; Kinghorn et al. 2015; Seleznev et al. 2006; Zhao et al. 2010).

PLAN is a rarer form of human neurodegeneration, caused by an autosomal recessive mutation in the *PLA2G6* (*Phospholipase A2 group VI*) gene (Gregory et al. 2026). *PLA2G6* encodes a calcium-independent phospholipase A2 enzyme (also called iPLA_2_-VIA and iPLA_2_β) (Banerjee et al. 2021b; Malley et al. 2018; Winstead et al. 2000). It is ubiquitously present in the cytoplasm, perinuclear membrane space, and mitochondria of mammalian tissues (Barbour and Ramanadham 2017; Ramanadham et al. 2004; Zhao et al. 2010; Zhou and Tian 2018), including in the neurons of the monkey brain (Ong et al. 2005). It participates in phospholipid remodeling and is essential for maintaining membrane integrity, lipid balance, and cell signaling through the Land’s cycle (Kurtovic-Kozaric et al. 2024). Phospholipid remodeling by PLA2G6 contributes to proper neuronal functions, as phospholipids contribute to vesicular movements, synaptic functions, and cell organelle integrity (Antonny et al. 2015). Additionally, it regulates cell proliferation, differentiation and death, inflammation, mitochondrial integrity, and fertility (Abi Nahed et al. 2016; Beck et al. 2011; Ramanadham et al. 2015; Turk et al. 2019). Loss of PLA2G6 function leads to abnormal lipid accumulation, enhanced lipid peroxidation, and mitochondrial dysfunction and degradation (Banerjee et al. 2021b; Kinghorn and Castillo-Quan 2016; Liu et al. 2024) that share significant molecular parallels with more prevalent neurodegenerative illnesses such as Alzheimer’s and Parkinson’s. Clinically, PLAN encompasses a spectrum of phenotypes, including infantile neuroaxonal dystrophy, atypical neuroaxonal dystrophy, and adult-onset dystonia-parkinsonism, often accompanied by brain iron accumulation (Beck et al. 2015; Kinghorn et al. 2015), promoting axonal dysfunction of the dopaminergic neurons with the progression of age (Beck et al. 2016). PLAN patients exhibit slow movement, impaired balance, muscle stiffness, and a shorter lifespan (Beck et al. 2016; Shinzawa et al. 2008). A robust treatment for PLAN is not available, mainly because the underlying mechanisms of this disease are still not fully known (Iliadi et al. 2018).

A major mechanism that ties the PLA2G6 mutation to mitochondrial abnormality needs detailed evaluation. PLA2G6 repairs mitochondrial membrane damage triggered by ROS-induced peroxidation, restores mitochondrial membrane potential, and inhibits apoptosis caused by mitochondrial damage (Seleznev et al. 2006; Zhao et al. 2010). Due to the *PLA2G6* mutation, mitochondrial membrane potential reduces, and mitochondrial Reactive Oxygen Species (ROS) increases in human fibroblast cells (Kinghorn et al. 2015). Additionally, the mitochondrial inner membrane degenerates in the proximal axons of the central and peripheral nervous systems of mice (Beck et al. 2011). Despite growing evidence that loss of PLA2G6 activity is associated with mitochondrial abnormalities, several key aspects of this relationship remain poorly defined. For example, a systematic analysis of how mitochondrial architecture changes across multiple tissues and over the course of aging in both sexes carrying the *PLA2G6* mutation is lacking. It is also unclear whether these structural alterations are accompanied by progressive loss of mitochondrial abundance and impairment of mitochondrial function in PLAN. In particular, the consequences of PLA2G6 deficiency for cellular bioenergetics and redox homeostasis have not been comprehensively evaluated in vivo. Furthermore, the transcriptional regulation of the genes that control mitochondrial production, fusion and fission dynamics, and mitophagy has also not been analyzed in PLAN. Thus, identifying these mechanisms across sexes, tissues, and ages remains essential for developing effective treatments of PLAN.

To address these questions, scientists have employed a *Drosophila melanogaster* (fruit flies) model of PLAN, carrying a homozygous null mutation in the iPLA_2_-VIA gene (Banerjee et al. 2021b; Iliadi et al. 2018; Kinghorn et al. 2015; Meimoun et al. 2025). Similar to its mammalian ortholog, PLA2G6, the *Drosophila* iPLA_2_-VIA protein contains highly conserved nine ankyrin domains, an oxyanion hole, a catalytic site, and a calmodulin-binding motif. Both proteins share over 50% of amino acid sequence similarity. Additionally, in flies, iPLA_2_-VIA is expressed in the mitochondria of the male and female germ cells, along with in other cell organelles in different tissues (Banerjee et al. 2021b; Liu et al. 2024). Compared with control flies, homozygous iPLA_2_-VIA null mutants exhibit age-dependent major climbing defects, severely reduced lifespan in both sexes, and female-specific fertility defects (Banerjee et al. 2021b; Kinghorn et al. 2015). The age-dependent locomotor decline in the mutant flies is associated with progressive loss of dopaminergic neurons (Mori et al. 2019). Furthermore, heads isolated from one-month-old mutant flies display pronounced mitochondrial structural abnormalities accompanied by reduced mitochondrial membrane potential, decreased ATP production, and reduced cellular respiration in the complexes I and II of the electron transport chain (Kinghorn et al. 2015). Similarly, ovaries from three-week-old mutant females show markedly reduced mitochondrial membrane potential and abnormal mitochondrial distribution in the nurse cells (Banerjee et al. 2021b). Together, these findings suggest a strong correlation between progressive mitochondrial dysfunction and the age-dependent neuromuscular pathology observed in PLAN flies. However, a systemic analysis of mitochondrial disruption in PLAN flies is still absent.

Therefore, we used PLAN flies to perform a comprehensive analysis of mitochondrial properties following loss of iPLA2-VIA. We examined neuronal (head), muscular (thorax), and reproductive (ovary) tissues from young (7-day-old) and old (3-week-old) flies of both sexes to characterize tissue-, age-, and sex-dependent mitochondrial dysregulation. Our analyses revealed that iPLA_2_-VIA mutant flies exhibit severe mitochondrial structural abnormalities, including disrupted mitochondrial membranes, degenerated cristae, abnormal mitochondrial morphology, and reduced mitochondrial abundance across multiple tissues. This reduction in mitochondrial number was further supported by a significant decrease in mitochondrial DNA content in whole adult mutant flies, while the amount of nuclear genome remained unchanged compared to controls. In addition, brains isolated from 7-day-old and 3-week-old male and female mutant flies showed reduced intensity for mitochondrial molecular markers. A clumpy mitochondrial phenotype was also observed in 3-week-old mutant female brain samples.

To determine whether these structural and quantitative mitochondrial defects were associated with functional impairment, we measured ATP and ROS levels. These assays were performed in the head, thorax, and ovary of 7-day-old and 3-week-old, as well as in whole adult control and iPLA_2_-VIA mutant flies. iPLA_2_-VIA mutant flies showed reduced ATP production in most tissues and at the whole-body level. They also showed age-, sex-, and tissue-dependent changes in ROS levels. *Sod2* expression was selectively reduced in young mutant females but remained unchanged in most other groups, suggesting that ROS alterations are not explained solely by broad suppression of antioxidant gene expression.

Because mitochondrial structure and function depend on coordinated maintenance pathways, we next analyzed the expression of genes involved in mitochondrial biogenesis, fusion and fission dynamics, and quality control. We first examined *mTOR* and *PGC-1α* expressions to determine whether the reduced mitochondrial number observed in iPLA_2_-VIA mutant flies resulted from inactivation of mitochondrial biogenesis pathways. Both *mTOR* and *PGC-1α* expressions were reduced in iPLA_2_-VIA mutant flies. Genes involved in mitochondrial fusion showed a selective pattern, in which *Mfn1* remained unchanged, *Mfn2* increased only in young mutant females, and *Opa1* was significantly reduced in young and old iPLA_2_-VIA mutants. Fission-related genes also changed mainly in young flies, with altered *Drp1* expression and reduced *Fis1* expression. In addition, *Pink1* showed age- and sex-dependent changes, suggesting altered mitochondrial quality-control signaling. *Trap1* was reduced in old mutants, whereas *Sirtuin 6* remained largely unchanged.

Together, this integrated approach reveals progressive, tissue-specific mitochondrial degeneration in a *Drosophila* model of PLA2G6-associated neurodegeneration. The combined structural, molecular, and functional findings suggest that loss of iPLA_2_-VIA disrupts mitochondrial membrane integrity, reduces mitochondrial abundance, impairs ATP production, and alters redox balance. These defects appear to be associated with impaired mitochondrial biogenesis, altered fusion and fission dynamics, and disrupted mitochondrial quality-control pathways. Overall, this study provides evidence that defective mitochondrial maintenance is a central feature of PLAN pathogenesis and may contribute to the progressive neuromuscular decline observed following loss of iPLA_2_-VIA.

## Materials and Methods

### Fly strain and husbandry

We used the previously described two *Drosophila melanogaster* fly strains.: t; l/Sm6; Δ11/Δ11 as the control strain and t; l/Sm6; Δ23/Δ23 as the homozygous null mutant (*iPLA_2_-VIA^Δ23^*) strain. Homozygous control and null mutant flies were used in all experiments. We used sex and age-matched young (7 days old) and old (3 weeks old) control and mutant adult flies. Flies were cultured and maintained in a Percival DR-36NL *Drosophila* fly incubator with a set temperature of 25 °C. All fly crosses were set at 25 °C. Flies were cultured in vials with freshly prepared food made from nuti-fly (Genesee Scientific 66-121). 20 ml 20% tegosept (Apex 20-258) and 20 ml propionic acid (Ward’s Science 470302-290) were added per 3.5 litres of food.

### Transmission Electron Microscopy (TEM)

Head, thorax and ovaries from flies (n=10 per sex, per genotype) at both 7 days and 3 weeks old were fixed in a solution of 2.5% glutaraldehyde (Thermo Scientific A17876.0F) and 2% paraformaldehyde (Thermo Scientific 043368.9M) prepared in 0.2 M Sorenson’s phosphate buffer (pH 7.3) (Electron Microscopy Sciences 1160005) for 1 hour at room temperature, followed by overnight fixation at 4 °C. Samples were rinsed thoroughly in 0.05 M cacodylate buffer and post-fixed with 1% osmium tetroxide (Sigma-Aldrich 251755) for 1 hour at room temperature. After three additional buffer washes, tissues were dehydrated through a graded ethanol series (25%, 50%, 75%, 85%, 95%, and 100%), followed by two washes in 100% acetone (Thermo Scientific L10407.0F). Infiltration was performed using increasing concentrations of epoxy resin (LX-112 Embedding Kit, Ladd Research) mixed with acetone (ratios of 4:1, 1:1, and 1:4), followed by two incubations in 100% resin. Samples were embedded in fresh resin and polymerized at 60 °C for 48 hours. Ultrathin sections (∼67 nm) were cut using an ultramicrotome (RMC powertome), stained with uranyl acetate (Electron Microscopy Sciences 22400) and lead citrate (Sigma-Aldrich 15326), and then imaged using a Hitachi H7650 transmission electron microscope at multiple magnifications: 3000x, 6000x, and 10000x (Hurd et al., 2015). Mitochondrial numbers were counted using the images taken at 3000x magnification.

### Mitotracker and Hoechst staining of brain samples

MitoTracker staining was performed using a protocol adapted from Wong et al. (2020). Adult brains from 7-day-old and 3-week-old male and female iso-control and iPLA_2_-VIA mutant flies were dissected in ice-cold 1× phosphate-buffered saline (PBS). MitoTracker Green FM (Ex/Em: 490/516 nm; Thermo Scientific, M7514) was prepared as a 1 mM stock solution by dissolving 50 μg of dye in 74.4 μL anhydrous dimethyl sulfoxide (DMSO). Hoechst 33342 (Ex/Em: 350/461 nm; Thermo Scientific) was reconstituted in deionized water to generate a 10 mg/mL stock solution, and a 1 mg/mL intermediate solution was prepared by dilution in 1× PBS (10X PBS:1.37 M NaCl, 27 mM KCl, 100 mM Na2HPO4, 18 mM KH2PO4, pH 7.4). For staining, brains were incubated in a combined dye solution containing MitoTracker Green FM at a final concentration of 200 nM, and Hoechst working solution was added at a 100:1 ratio. All dye solutions and samples were protected from light throughout the procedure to minimize photobleaching. All dye solutions and samples were protected from light throughout the procedure. Dissected brains were washed in 1× PBS and incubated in pre-warmed dye solution at 37 °C for 45 min at room temperature, in the dark. Following incubation, brains were washed twice in 1× PBS, fixed in 4% paraformaldehyde (PFA) (Thermo Scientific 043368.9M) for 5 min, washed again in 1× PBS, and mounted in 80% antifade glycerol under sealed coverslips (Wong et al. 2020).

Bright-field and Z-stack confocal fluorescence images were acquired using a Leica DMi8 Inverted THUNDER confocal spinning disk microscope at 40× magnification. Hoechst 33342 was detected using the 405 nm laser channel, and MitoTracker Green was detected using the 470 nm laser channel. MitoTracker Green FM was used to visualize mitochondria, while Hoechst 33342 was used to label nuclear DNA. Laser power and exposure settings were kept consistent between control and mutant samples within each experimental comparison.

### ATP Quantification

ATP levels were quantified using a luciferase-based chemiluminescence assay following the manufacturer’s protocol (ATP Determination Kit, Molecular Probes A22066). Adult fly heads, thoraxes, and ovaries (n=15 per sex, per genotype, per biological replicate with three biological replicates), or whole adult male and female flies (n=5 per sex, per genotype, per biological replicate with six biological replicates) at both 7 days and 3 weeks old were dissected. Fresh tissues were rapidly homogenized on ice in 100 μL extraction buffer (6 M guanidine-HCl, 100 mM Tris, 4 mM EDTA, pH 7.8) using a pellet pestle. An aliquot (20 μL) of the homogenate was reserved for protein quantification, while the remaining homogenate was incubated at 95 °C for 5 min, followed by centrifugation at 13,000 rpm (Labnet Prism R Refrigerated Micro-Centrifuge, C2500 R) for 3 min at 4 °C. The supernatant was collected and diluted with dilution buffer (100 mM Tris, 4 mM EDTA, pH 7.8). ATP standards were prepared by serial dilution of the supplied ATP stock. For the assay, 10 μL of standards or diluted samples were loaded into a white opaque 96-well plate, and the reaction was initiated by adding 100 μL of freshly prepared ATP reaction mix. Luminescence was measured immediately using a plate reader (BioTek Synergy H1), and the average of three sequential readings was taken for each well. ATP concentrations were calculated from the standard curve and normalized to the total protein content, which was determined separately by the Bradford assay (Tennessen et al. 2014). Final ATP levels were expressed as relative light units per microgram of protein (RLU/μg protein).

### ROS Quantification

Intracellular reactive oxygen species (ROS) levels were measured using a fluorescence-based assay (ROS Assay Kit, Abcam ab238535) as described in (Arzoo et al. 2025). Adult fly heads, thoraxes, and ovaries (n = 15 per sex per genotype per biological replicate, with three biological replicates) at both 7 days and 3 weeks old were dissected and homogenized in ice-cold 1X PBS. Homogenates were centrifuged at 10,000 × g (Labnet Prism R Refrigerated Micro-Centrifuge, C2500 R) for 5 min at 4 °C, and the supernatant was collected for analysis. The supernatant was diluted with 1X PBS. Hydrogen peroxide (H₂O₂) standards were freshly prepared by serial dilution. For the assay, 50 μL of standards or diluted samples were loaded into a black 96-well plate. Samples and standards were incubated with the supplied catalyst, followed by the addition of the DCFH working solution. The plate was protected from light and incubated at room temperature for 30 minutes. Fluorescence was measured using a plate reader (BioTek Synergy H1) with an excitation wavelength of 480 nm and an emission wavelength of 530 nm. ROS levels were calculated from the H₂O₂ standard curve and normalized to the corresponding protein concentration of each sample, as determined separately by the Bradford assay. Final ROS values were expressed as fluorescence units per microgram of protein (Arzoo et al., 2025).

### Bradford Assay

For normalization of our ATP and ROS data, we performed the Bradford assay to find the total protein concentration of the samples (Banerjee et al. 2021a; Tennessen et al. 2014). An aliquot of each tissue homogenate was centrifuged to remove insoluble debris, and the supernatant was used for protein determination. A BSA standard curve (0–2 mg/mL) was prepared by serial dilution of BSA in the dilution buffer or PBS. Five microliters of standards or samples were loaded in duplicate into a clear 96-well plate, followed by the addition of 200 μL Bradford reagent (BioRad 5000205). Plates were incubated at room temperature for 30 min, and absorbance was measured at 595 nm using a plate reader (BioTek Synergy H1). Protein concentrations were calculated from the standard curve and used to normalize ATP and ROS measurements.

### Quantitative Real-Time PCR (RT-qPCR) Analysis

Total RNA was isolated from whole flies (20 flies per sample) at 7 days and 3 weeks of age using TRI reagent (Sigma T9424). RNA quality was verified by A260/280 and A260/230 ratio, and by running 2 µg of RNA in 1% Agarose gel to check for RNA degradation. Complementary DNA (cDNA) was synthesized from 1 µg RNA using the iScript Reverse Transcription Supermix for RT-qPCR (Bio-Rad 1708841) according to the manufacturer’s protocol. Primer sequences (FP = Forward Primer, and RP = Reverse Primer), listed in Table 1, were designed using the Integrated DNA Technologies Primerquest qPCR primer design tool (Kalendar 2025) or previously published articles. They were validated for amplification efficiency and specificity. Quantitative PCR (10 µL) was performed using SsoFast EvaGreen Supermix (Bio-Rad, 172-5201) on a Bio-Rad MiniOpticon Real-Time PCR System. Expression levels of target genes were normalized to the endogenous reference gene *RpL32’s* expression within the same samples (Banerjee et al. 2021a). Relative gene expression levels were calculated using the 2^⁻ΔΔCT^ method (Bustin et al. 2009). All RT-qPCR experiments were conducted in accordance with the MIQE guidelines (Banerjee et al. 2021a; Bustin et al. 2009).

**Table 1.**
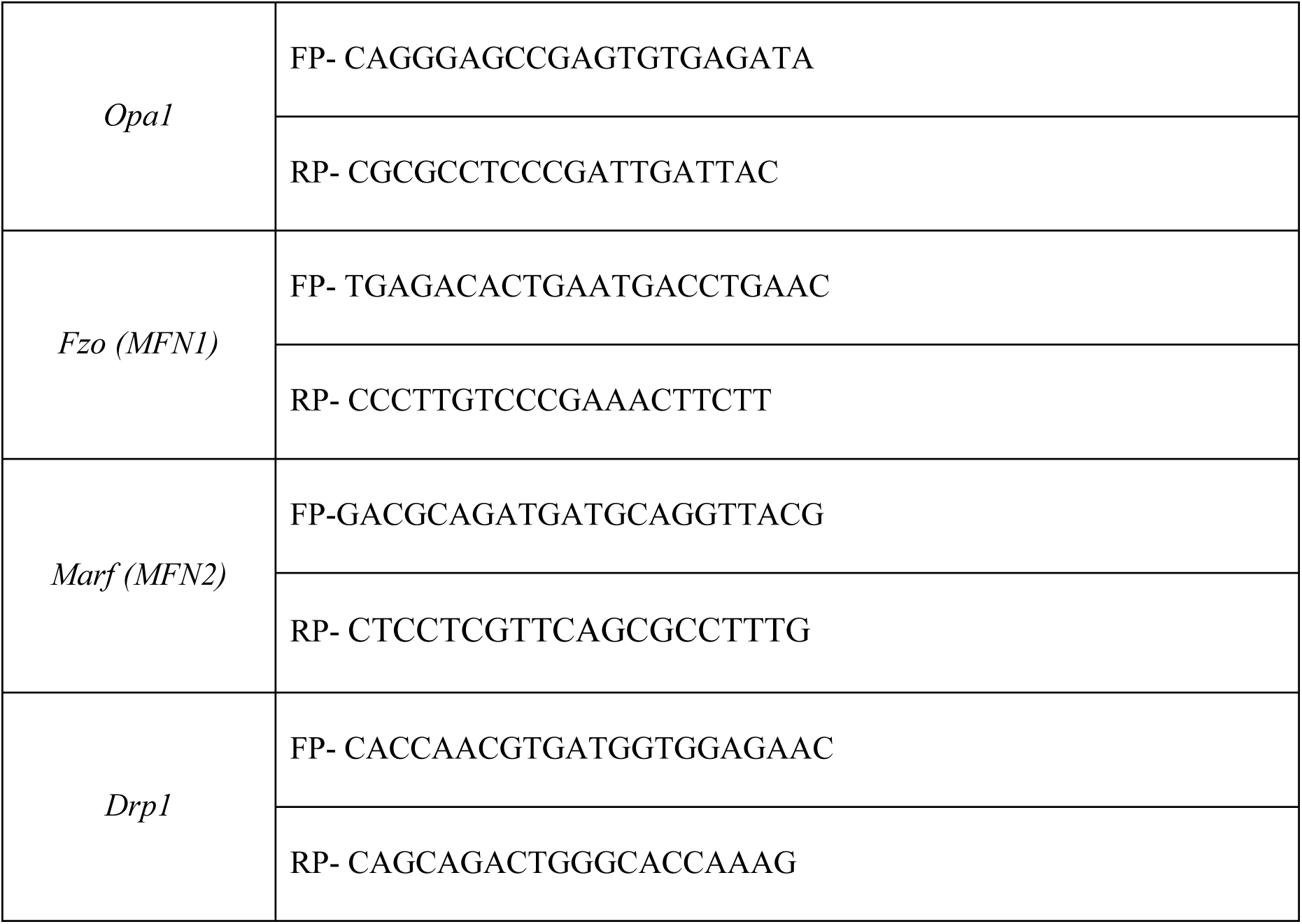
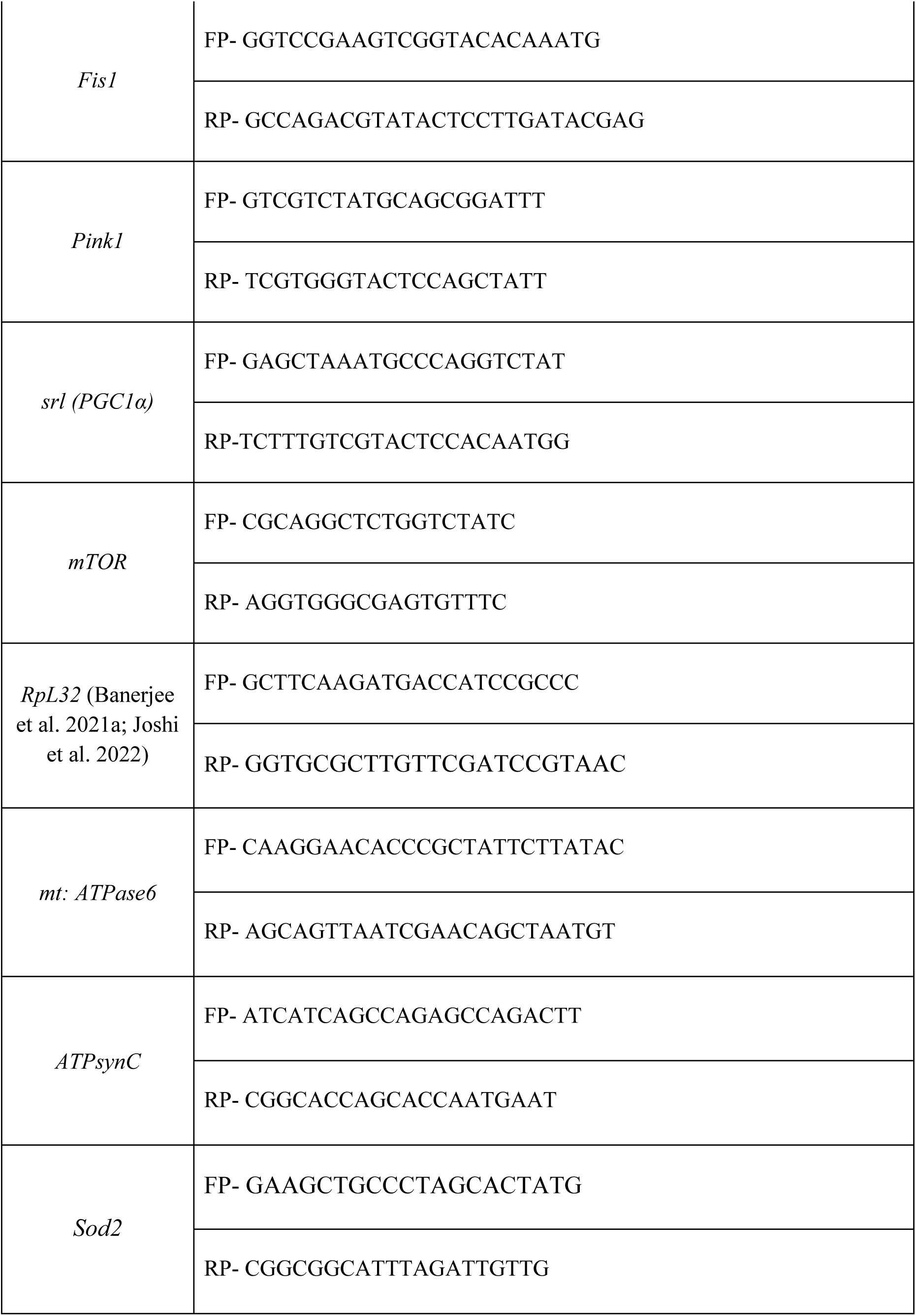
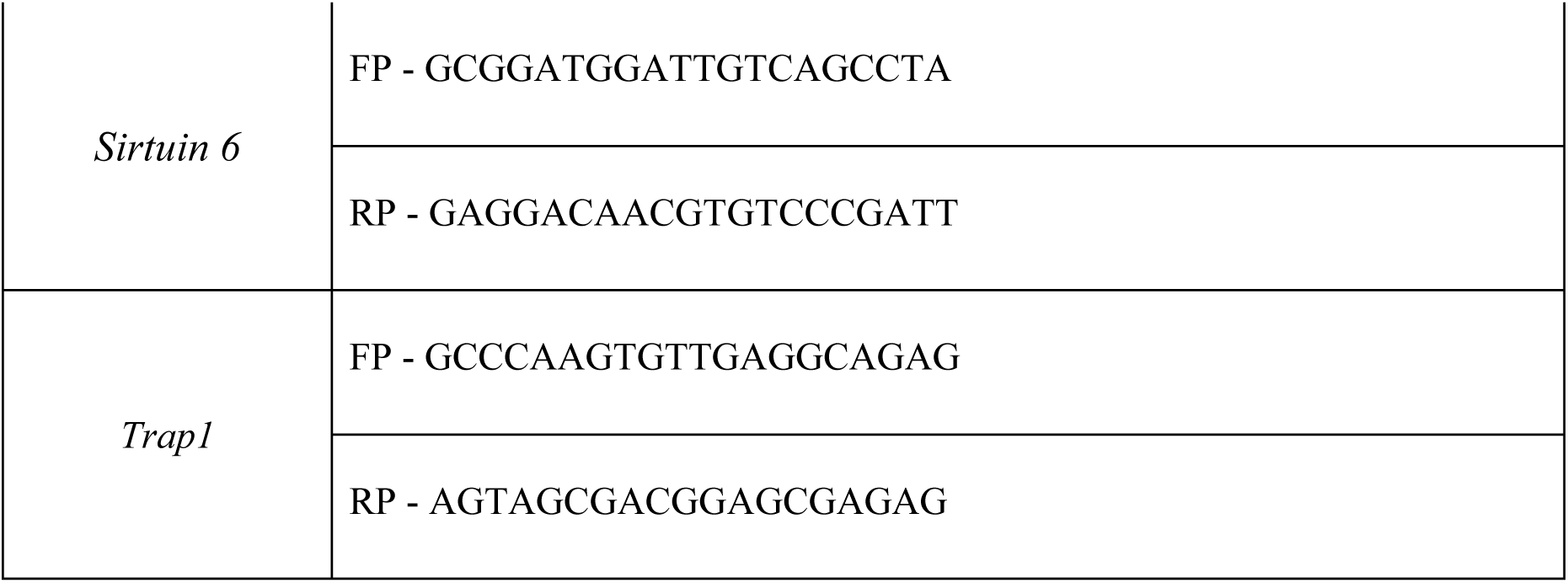
List of the forward and reverse primer sequences used for RT-qPCT.

### Quantification of Mitochondrial DNA

Total DNA was extracted from 10 whole adult *Drosophila melanogaster*. Flies were homogenized in a lysis buffer (Tris-HCl, EDTA, SDS, and proteinase K) and incubated at 37 °C for 60 min to facilitate protein digestion. The lysate was then incubated at 95 °C for 5 min to inactivate proteinase K. Following centrifugation, the supernatant containing DNA was collected, and DNA was precipitated using isopropanol, washed with 70% ethanol, and resuspended in DEPC water. Mitochondrial DNA content was quantified relative to nuclear DNA by quantitative real-time PCR using the ratio of the mitochondrial gene *ATPase6* to the nuclear gene *ATPSynC* (Rera et al. 2011).

### Statistical Analysis

All statistical analyses were performed using GraphPad Prism software. Comparisons between iso-control and iPLA_2_-VIAΔ23 mutant groups within the same age and sex were performed using an unpaired two-tailed Student’s *t*-test. Data are presented as mean ± SEM unless otherwise indicated. Statistical significance was defined as *p* < 0.05. Significance levels are indicated as follows: ns, not significant; *p* < 0.05; \**p* < 0.01; \*\**p* < 0.001; and \*\*\**p* < 0.0001.

## Results

### 1. Loss of iPLA_2_-VIA causes mitochondrial ultrastructural abnormalities across tissues

To determine whether loss of iPLA_2_-VIA affects mitochondrial integrity in vivo, we examined mitochondrial ultrastructure in the brains, thoraces, and ovaries of control and iPLA_2_-VIA *mutant Drosophila melanogaster* using transmission electron microscopy (TEM). Analyses were performed in both young (7-day-old) and aged (3-week-old) male and female flies to assess age-and sex-dependent effects (Figures 1).

**Figure 1:**
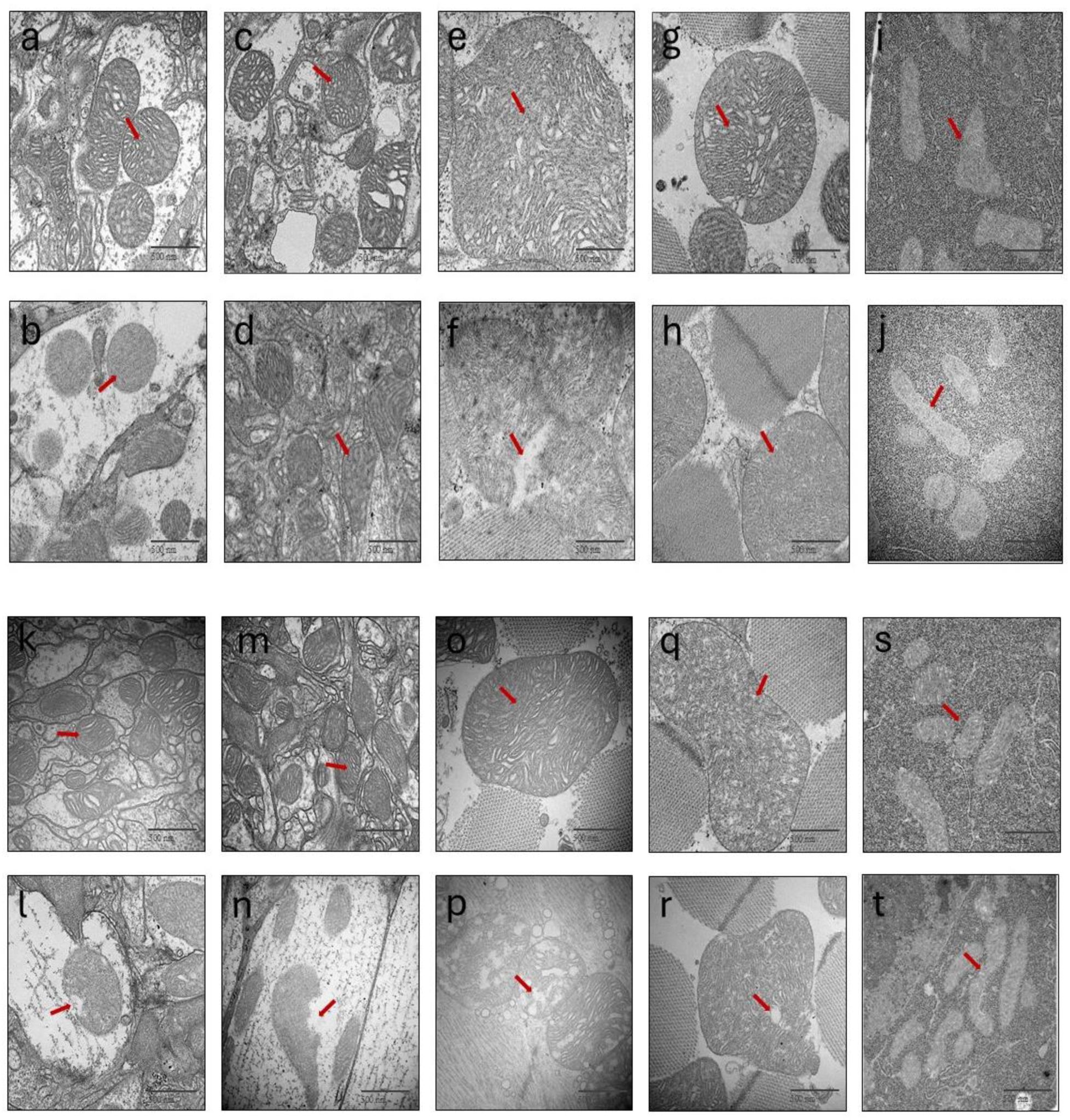
Representative transmission electron microscopy (TEM) images showing mitochondrial ultrastructure in iso-control and iPLA_2_-VIA mutant Drosophila melanogaster tissues at 7 days and 3 weeks of age. Panels A–J show tissues from 7-day-old flies, and panels K–T show tissues from 3-week-old flies. For 7-day-old flies, panels A and B show female brain, C and D show male brain, E and F show female thorax, G and H show male thorax, and I and J show ovary. For 3-week-old flies, panels K and L show female brain, M and N show male brain, O and P show female thorax, Q and R show male thorax, and S and T show ovary. Scale bar = 500 nm.

In 7-day-old flies, iso-control tissues showed mitochondria with relatively well-preserved morphology and organized internal cristae. In the 7-day-old female brain, control mitochondria showed clear mitochondrial profiles with visible cristae structure (Figure 1A). In contrast, mitochondria in the 7-day-old mutant female brain showed altered morphology and disrupted internal organization, with less clearly defined cristae structure (Figure 1B). A similar pattern was observed in male brains. Control male brain mitochondria showed organized cristae and preserved mitochondrial architecture (Figure 1C), whereas mutant male brain mitochondria displayed abnormal internal structure and disrupted cristae organization (Figure 1D). These observations indicate that mitochondrial ultrastructural abnormalities are already present in the nervous tissue of young adult iPLA_2_-VIA mutant flies.

Mitochondrial defects were also observed in thoracic tissue from young flies. In the 7-day-old iso-control female thorax, mitochondria showed dense and organized cristae structure (Figure 1E). However, mitochondria from the 7-day-old mutant female thorax showed disrupted internal architecture and poorly organized cristae (Figure 1F). In the 7-day-old male thorax, control mitochondria maintained a defined shape with visible cristae (Figure 1G), while mutant mitochondria showed abnormal morphology and reduced internal organization (Figure 1H). These findings suggest that loss of iPLA_2_-VIA affects mitochondrial structure not only in brain tissue but also in metabolically active muscle tissue.

We next examined mitochondrial morphology in the ovary of young flies. In 7-day-old iso-control ovaries, mitochondria showed well-defined structure and organized internal membranes (Figure 1I). In contrast, mitochondria in 7-day-old iPLA_2_-VIA mutant ovaries showed abnormal morphology, disrupted cristae organization, and altered internal architecture (Figure 1J). This indicates that mitochondrial structural defects are also present in reproductive tissue at an early adult stage.

Mitochondrial abnormalities became more evident in aged flies. In 3-week-old iso-control female brains, mitochondria retained a well-organized structure with visible cristae (Figure 1K). In contrast, 3-week-old mutant female brains showed mitochondria with disrupted morphology, reduced cristae organization, and compromised internal structure (Figure 1L). In aged male brains, iso-control mitochondria also showed relatively preserved internal organization (Figure 1M), whereas mutant mitochondria showed abnormal structure and loss of normal cristae architecture (Figure 1N). These observations suggest that mitochondrial defects persist and may worsen with age in the neural tissue of iPLA_2_-VIA mutants.

Aged thoracic tissues also showed clear mitochondrial abnormalities in mutant flies. In the 3-week-old iso-control female thorax, mitochondria showed a defined shape and organized cristae pattern (Figure 1O). In contrast, mitochondria from 3-week-old mutant female thorax showed disrupted cristae, irregular morphology, and less organized internal structure (Figure 1P). Similarly, the 3-week-old iso-control male thorax showed mitochondria with preserved internal architecture (Figure 1Q), whereas the mutant male thorax showed mitochondria with abnormal morphology and disrupted cristae organization (Figure 1R). These findings indicate that mitochondrial structural defects in the thorax are maintained in aged mutant flies.

Finally, aged ovarian tissue also showed mitochondrial abnormalities in iPLA_2_-VIA mutants. In 3-week-old iso-control ovaries, mitochondria displayed visible internal organization and preserved morphology (Figure 1S). In contrast, mitochondria from 3-week-old mutant ovaries showed disrupted internal architecture and altered cristae organization (Figure 1T). The presence of mitochondrial abnormalities in aged ovaries supports the idea that loss of iPLA_2_-VIA affects mitochondrial structure across multiple tissue types.

Together, these TEM data show that loss of iPLA_2_-VIA causes widespread mitochondrial ultrastructural defects in brain, thorax, and ovary. These abnormalities were observed in both male and female flies and were present as early as 7 days of age. In aged flies, mitochondrial defects remained evident across tissues, suggesting a progressive or persistent impairment of mitochondrial structural maintenance. The observed defects, including disrupted cristae organization, abnormal mitochondrial morphology, and compromised internal architecture, support the conclusion that iPLA_2_-VIA is required for maintaining mitochondrial ultrastructure in adult *Drosophila* tissues.

### 2. iPLA_2_-VIA Mutants Exhibit a Significant Reduction in Mitochondrial Number

Because TEM images revealed widespread mitochondrial ultrastructural abnormalities in iPLA_2_-VIA mutant flies, we next quantified mitochondrial number to determine whether loss of iPLA_2_-VIA also affects mitochondrial abundance. Mitochondrial counts were performed from TEM images of brain, thorax, and ovary tissues from 7-day-old and 3-week-old iso-control and iPLA_2_-VIA mutant flies. Comparisons were made between control and mutant groups within the same sex, age, and tissue.

In 7-day-old flies, mitochondrial number was significantly reduced in the brains of both female and male iPLA_2_-VIA mutants compared with age-matched iso-controls. In 7-day-old female brains, mutant flies showed a clear decrease in mitochondrial number relative to controls (Figure 2A). A similar reduction was observed in 7-day-old male brains, where iPLA_2_-VIA mutants also had significantly fewer mitochondria than iso-control males (Figure 2B). These findings indicate that mitochondrial depletion is already present in young adult mutant neural tissue in both sexes.

**Figure 2:**
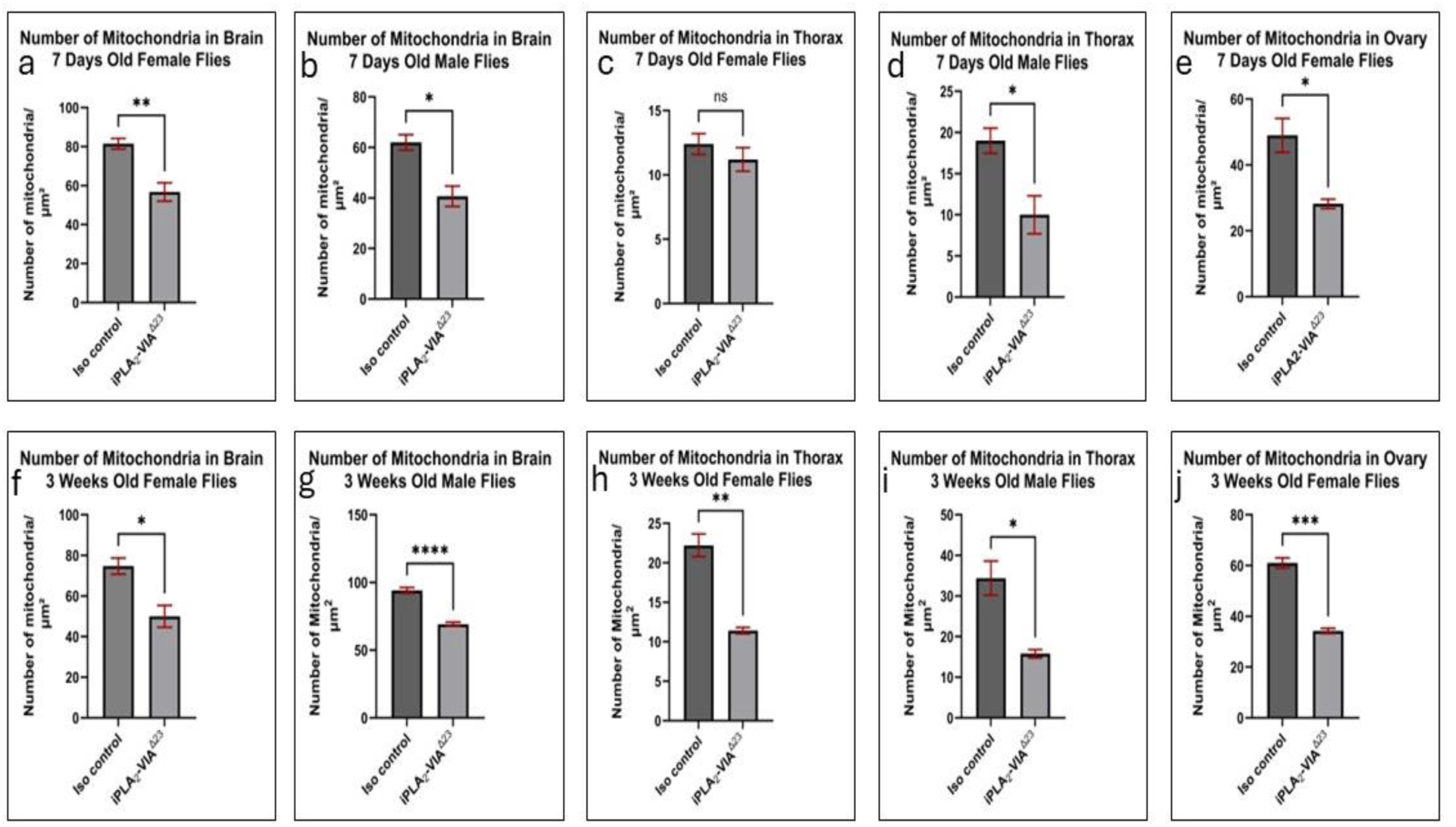
Quantification of mitochondrial number from transmission electron microscopy images of iso-control and iPLA_2_-VIA mutant Drosophila melanogaster tissues at 7 days and 3 weeks of age. Panels A–E show mitochondrial counts from 7-day-old flies: female brain (A), male brain (B), female thorax (C), male thorax (D), and female ovary (E). Panels F–J show mitochondrial counts from 3-week-old flies: female brain (F), male brain (G), female thorax (H), male thorax (I), and female ovary (J). Mitochondrial number was reduced in mutant brains of both sexes at 7 days and 3 weeks, with additional reductions observed in male thorax and ovary at 7 days and in all examined tissues at 3 weeks. Data are presented as mean ± SEM. Statistical comparisons were performed between iso-control and iPLA_2_-VIA mutant groups within the same age, sex, and tissue using an unpaired two-tailed t-test.

In thoracic tissue from 7-day-old flies, the effect of iPLA_2_-VIA loss was more sex-dependent. In the 7-day-old female thorax, mitochondrial number was slightly reduced in mutants compared with controls, but this difference was not statistically significant (Figure 2C). In contrast, the 7-day-old male mutant thorax showed a significant reduction in mitochondrial number compared with the iso-control male thorax (Figure 2D). This suggests that mitochondrial loss in young thoracic tissue is more pronounced in males than in females at this early adult stage.

Mitochondrial number was also reduced in reproductive tissue from young flies. In 7-day-old female ovaries, iPLA_2_-VIA mutants showed significantly fewer mitochondria compared with iso-control ovaries (Figure 2E). This finding indicates that mitochondrial depletion is not restricted to neural or muscular tissues but is also present in reproductive tissue during early adulthood.

The reduction in mitochondrial number became more widespread in aged flies. In 3-week-old female brains, the mitochondrial number was significantly decreased in iPLA_2_-VIA mutants compared with iso-controls (Figure 2F). This reduction was even more pronounced in 3-week-old male brains, where mutant flies showed a highly significant decrease in mitochondrial number relative to controls (Figure 2G). These data show that mitochondrial depletion persists with age in the brain and is evident in both sexes.

Aged thoracic tissues also showed significant mitochondrial loss in mutant flies. In the 3-week-old female thorax, mitochondrial number was significantly reduced in iPLA_2_-VIA mutants compared with iso-control females (Figure 2H). Similarly, the 3-week-old mutant male thorax showed a significant reduction in mitochondrial number compared with age-matched controls (Figure 2I). Unlike the 7-day-old thorax, where only males showed a significant reduction, both male and female thoracic tissues were affected at 3 weeks of age. This suggests that mitochondrial loss in muscle tissue becomes more evident with aging.

Finally, mitochondrial number was significantly reduced in the ovaries of 3-week-old iPLA_2_-VIA mutant females compared with iso-control females (Figure 2J). The reduction observed in aged ovaries, together with the significant decrease already present at 7 days, indicates that reproductive tissue is also vulnerable to mitochondrial depletion following loss of iPLA_2_-VIA.

Overall, mitochondrial quantification showed that iPLA_2_-VIA mutant flies have reduced mitochondrial abundance across multiple tissues. At 7 days, significant reductions were observed in the female brain, male brain, male thorax, and ovary, while the female thorax showed no significant difference. By 3 weeks of age, mitochondrial number was significantly reduced in all examined tissues, including the female brain, male brain, female thorax, male thorax, and ovary. These data demonstrate that loss of iPLA_2_-VIA causes tissue- and sex-dependent mitochondrial depletion that becomes more widespread with age. Together with the ultrastructural defects shown in Figure 1, these findings indicate that iPLA_2_-VIA is required not only for maintaining mitochondrial structure but also for preserving mitochondrial abundance in adult fly tissues.

### 3. Fluorescence Imaging Shows Abnormal Mitochondrial Distribution in Old iPLA_2_-VIA Mutant Brains

To further examine mitochondrial distribution in young adult brains, we performed MitoTracker Green and Hoechst staining in 7-day-old iso-control and iPLA_2_-VIA mutant female and male flies. Z-stack confocal images were acquired at 40× magnification. MitoTracker Green was used to visualize mitochondria, while Hoechst staining was used to label nuclear DNA and show the overall organization of the brain tissue.

In 7-day-old iso-control female brains, MitoTracker Green staining was clearly visible across the brain tissue (Figure 3b). The merged image showed strong green mitochondrial staining together with Hoechst nuclear staining, confirming mitochondrial signal within the brain structure (Figure 3d). In contrast, 7-day-old iPLA_2_-VIA mutant female brains showed weaker MitoTracker Green staining compared with iso-control females (Figure 3f). The merged image also showed reduced green fluorescence in the mutant female brain, while Hoechst staining confirmed the presence of brain tissue (Figure 3h). These observations indicate that mitochondrial staining is reduced in young mutant female brains.

**Figure 3:**
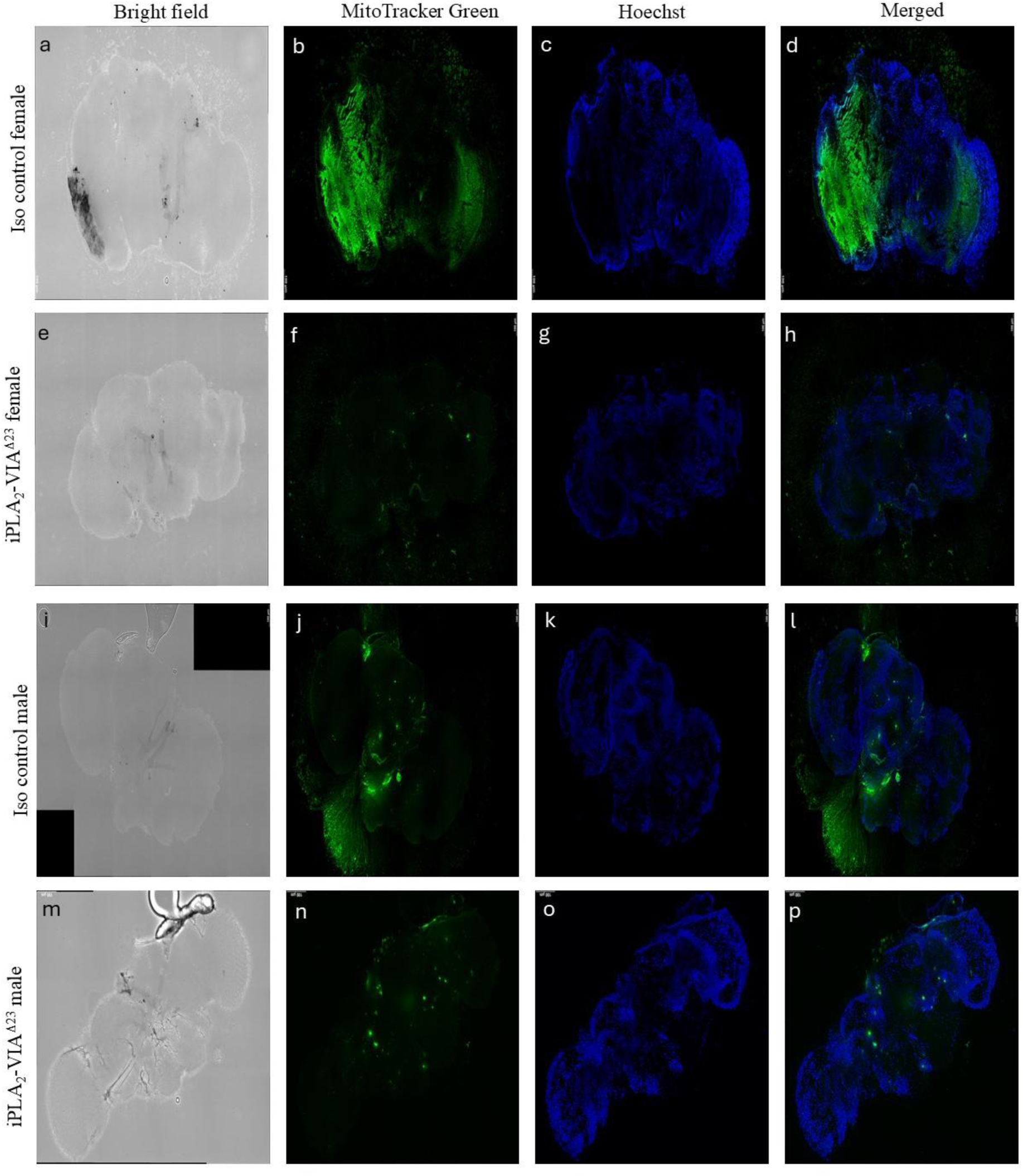
Z-stack confocal fluorescence images of brains from 7-day-old iso-control and iPLA_2_-VIA mutant flies. Images were acquired at 40× magnification. Panels a–d show the iso-control female brain, panels e–h show the iPLA_2_-VIA mutant female brain, panels i–l show the iso-control male brain, and panels m–p show the iPLA_2_-VIA mutant male brain. Columns show bright-field images, MitoTracker Green staining, Hoechst nuclear staining, and merged fluorescence images. MitoTracker Green was used to label mitochondria, and Hoechst was used to label nuclear DNA. Scale bars represent 100 μm.

A similar pattern was observed in 7-day-old male brains. Iso-control male brains showed detectable MitoTracker Green staining across the brain tissue (Figure 3j), with visible mitochondrial fluorescence in the merged image (Figure 3l). In contrast, 7-day-old iPLA_2_-VIA mutant male brains showed reduced and less broadly distributed MitoTracker Green staining (Figure 3n). The merged image of the mutant male brain also showed weaker green fluorescence compared with the iso-control male brain, while Hoechst staining remained visible (Figure 3p).

Together, these images show that MitoTracker Green staining is reduced in 7-day-old iPLA_2_-VIAΔ23 mutant brains compared with iso-control brains in both females and males. The reduction in mitochondrial staining in young adult mutant brains is consistent with the TEM-based mitochondrial quantification, which showed reduced mitochondrial number in 7-day-old mutant brain tissue. These data suggest that mitochondrial alterations are already detectable in neural tissue early in adulthood following loss of iPLA_2_-VIA.

To examine whether mitochondrial distribution worsens with age, we performed MitoTracker Green and Hoechst staining in 3-week-old iso-control and iPLA_2_-VIA mutant female and male flies. Z-stack confocal images were acquired at 40× magnification. MitoTracker Green was used to visualize mitochondria, while Hoechst staining was used to label nuclear DNA and show the overall structure of the brain tissue.

In 3-week-old iso-control female brains, MitoTracker Green staining was clearly detected across the brain tissue (Figure 4b). The merged image showed visible green mitochondrial staining together with Hoechst nuclear staining (Figure 4d). In contrast, 3-week-old iPLA_2_-VIA mutant female brains showed reduced and clumped MitoTracker Green staining compared with iso-control females (Figure 4f). In the merged image, the mitochondrial signal appeared more limited and punctate in the mutant female brain, while Hoechst staining confirmed the presence of brain tissue (Figure 4h).

**Figure 4:**
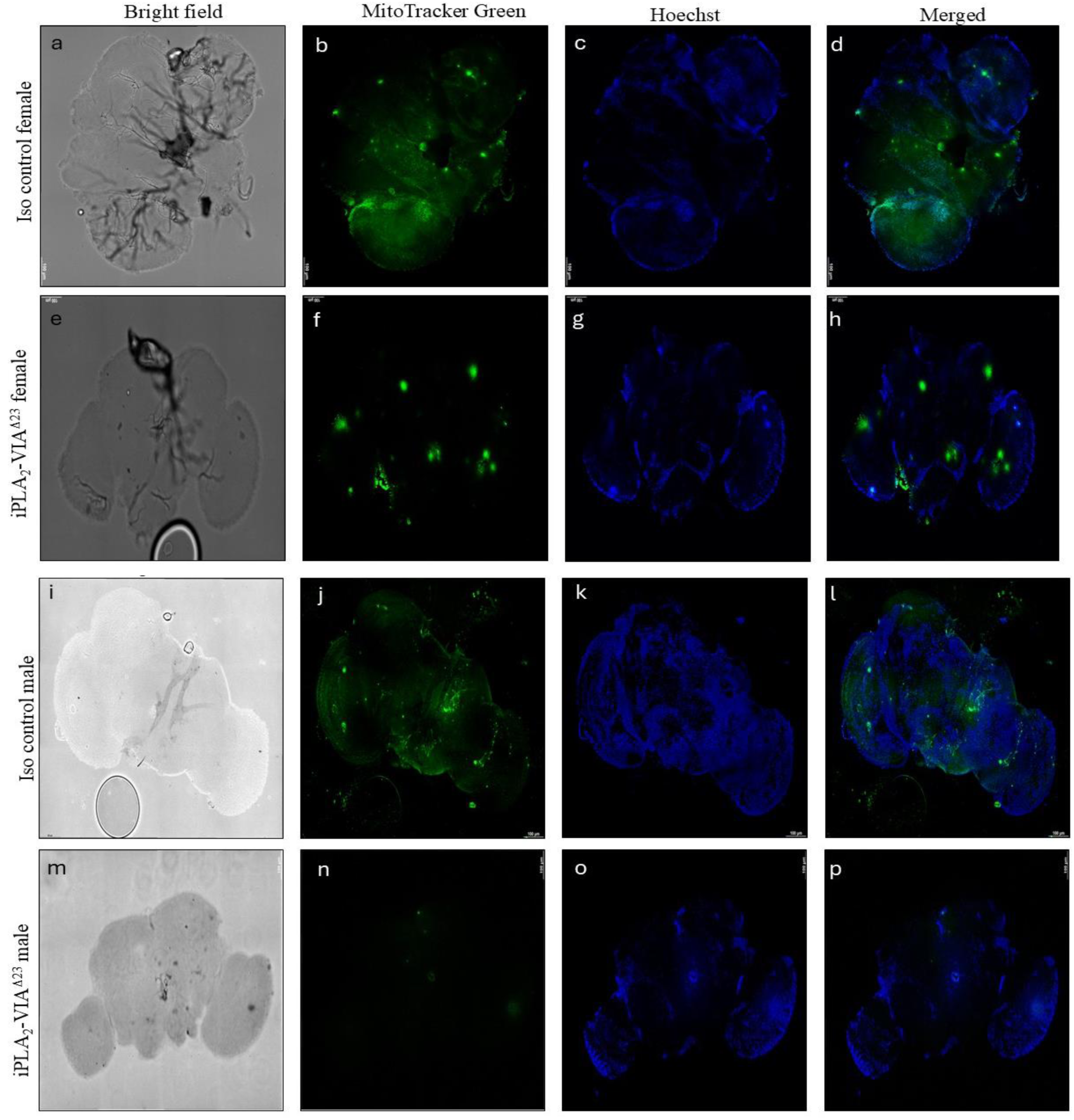
Z-stack confocal fluorescence images of brains from 3-week-old iso-control and iPLA_2_-VIA mutant *Drosophila melanogaster*. Images were acquired at 40× magnification. Panels a–d show the iso-control female brain, panels e–h show the iPLA_2_-VIA mutant female brain, panels i–l show the iso-control male brain, and panels m–p show the iPLA_2_-VIA mutant male brain. Columns show bright-field images, MitoTracker Green staining, Hoechst nuclear staining, and merged fluorescence images. MitoTracker Green was used to label mitochondria, and Hoechst was used to label nuclear DNA. Scale bars represent 100 μm.

A similar reduction was observed in aged male brains. In 3-week-old iso-control male brains, MitoTracker Green staining was broadly visible across the brain tissue (Figure 4j), and the merged image showed clear mitochondrial fluorescence within the Hoechst-stained brain structure (Figure 4l). In contrast, 3-week-old iPLA_2_-VIA mutant male brains showed markedly reduced MitoTracker Green staining (Figure 4n). The merged image also showed weaker mitochondrial fluorescence in the mutant male brain compared with the iso-control male brain (Figure 4p).

Together, these images show that aged iPLA_2_-VIA mutant brains have reduced MitoTracker Green staining compared with age-matched iso-control brains in both females and males. The reduction was visible in the MitoTracker channel and in the merged images, while Hoechst staining confirmed the presence of brain tissue in both control and mutant samples. These findings are consistent with the mitochondrial depletion observed by TEM-based quantification in 3-week-old mutant brains and support the conclusion that loss of iPLA_2_-VIA is associated with reduced mitochondrial abundance and altered mitochondrial staining patterns in aged neural tissue.

### 4. Reduced Mitochondrial Number is Confirmed by Reduction in Mitochondrial DNA Content in the Whole Body

Since TEM-based mitochondrial quantification showed reduced mitochondrial number in iPLA_2_-VIA mutant tissues, we next asked whether this reduction was also reflected at the level of mitochondrial DNA content. To test this, we performed qPCR using genomic DNA isolated from whole adult iso-control and iPLA_2_-VIA mutant flies. The mitochondrial DNA-encoded gene *ATPase6* was used as a marker of mitochondrial DNA abundance, while the nuclear DNA-encoded gene *ATPSynC* was used as a nuclear genomic control.

In 7-day-old female flies, *ATPase6* levels were significantly reduced in iPLA_2_-VIA mutants compared with iso-control females (Figure 5a). A similar reduction was observed in 7-day-old males, where mutant flies showed significantly lower *ATPase6* levels than iso-control males (Figure 5b). These findings indicate that mitochondrial DNA content is already reduced in young adult mutant flies of both sexes.

**Figure 5:**
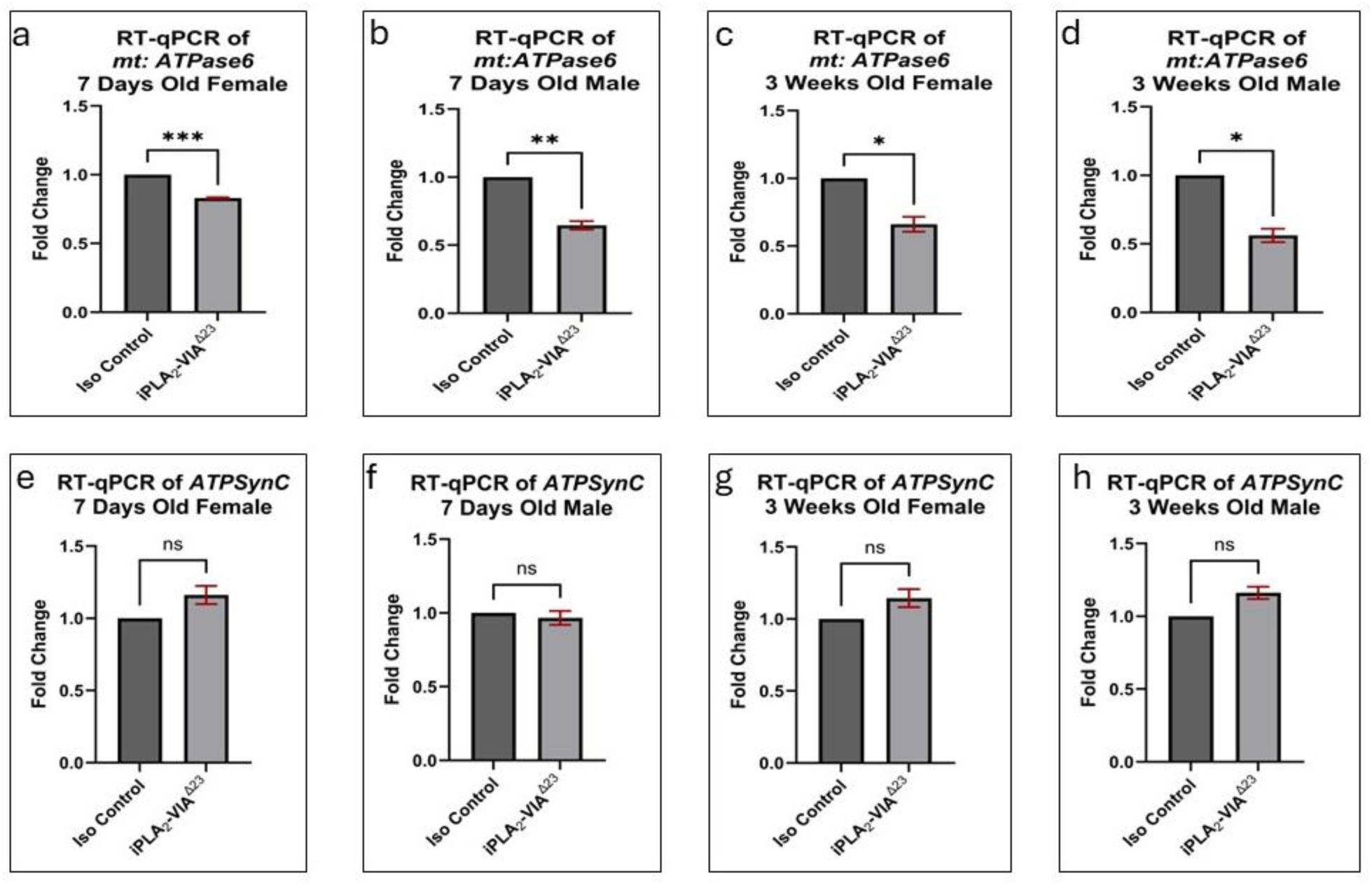
Quantification of mitochondrial DNA content by qPCR using genomic DNA isolated from iso-control and iPLA_2_-VIA mutant *Drosophila melanogaster*. The mitochondrial DNA-encoded gene *ATPase6* was used as a marker of mitochondrial DNA content, while the nuclear DNA-encoded gene *ATPSynC* was used as a nuclear genomic control. Panels A–D show *ATPase6* levels in 7-day-old females (a), 7-day-old males (b), 3-week-old females (c), and 3-week-old males (d). Panels E–H show *ATPSynC* levels in 7-day-old females (e), 7-day-old males (f), 3-week-old females (g), and 3-week-old males (h). *ATPase6* levels were significantly reduced in iPLA_2_-VIA mutants across both ages and sexes, whereas *ATPSynC* levels were not significantly changed. Data are presented as mean ± SEM. Statistical comparisons were performed between iso-control and iPLA_2_-VIA mutant groups within the same age and sex using an unpaired two-tailed *t*-test.

Reduced *ATPase6* levels were also observed in old mutant flies. In 3-week-old females, iPLA_2_-VIA mutants showed significantly decreased *ATPase6* levels compared with age-matched iso-control females (Figure 5c). Similarly, 3-week-old mutant males showed a significant reduction in *ATPase6* levels compared with iso-control males (Figure 5d). Thus, the reduction in mitochondrial DNA content was present at both young and aged stages and was observed in both female and male mutant flies.

To determine whether this reduction was specific to mitochondrial DNA rather than a general decrease in genomic DNA, we also measured the nuclear DNA-encoded gene *ATPSynC*. In contrast to *ATPase6*, *ATPSynC* levels were not significantly different between iso-control and iPLA_2_-VIA mutant flies in 7-day-old females (Figure 5e), 7-day-old males (Figure 5f), 3-week-old females (Figure 5g), or 3-week-old males (Figure 5h). The preservation of *ATPSynC* levels indicates that nuclear genomic DNA was not broadly reduced in the mutant samples.

These results show that iPLA_2_-VIA mutant flies have a selective reduction in mitochondrial DNA content, as shown by decreased *ATPase6* levels with unchanged *ATPSynC* levels. This molecular evidence supports the TEM-based observation of reduced mitochondrial abundance and suggests that loss of iPLA_2_-VIA is associated with mitochondrial depletion at both the structural and genetic levels.

### 5. Mitochondrial Dysfunction in iPLA_2_-VIA Mutants Leads to Reduced ATP and ROS Levels

To determine the functional consequences of mitochondrial ultrastructural defects, reduced mitochondrial number, and decreased mitochondrial DNA content, we next examined whether these changes were associated with altered mitochondrial function. ATP production and ROS levels were measured in heads, thoraxes, and ovaries from 7-day-old and 3-week-old iso-control and iPLA_2_-VIA mutant flies. ATP was used as a readout of mitochondrial bioenergetic output, while ROS was measured to assess changes in redox homeostasis (Figure 6).

**Figure 6:**
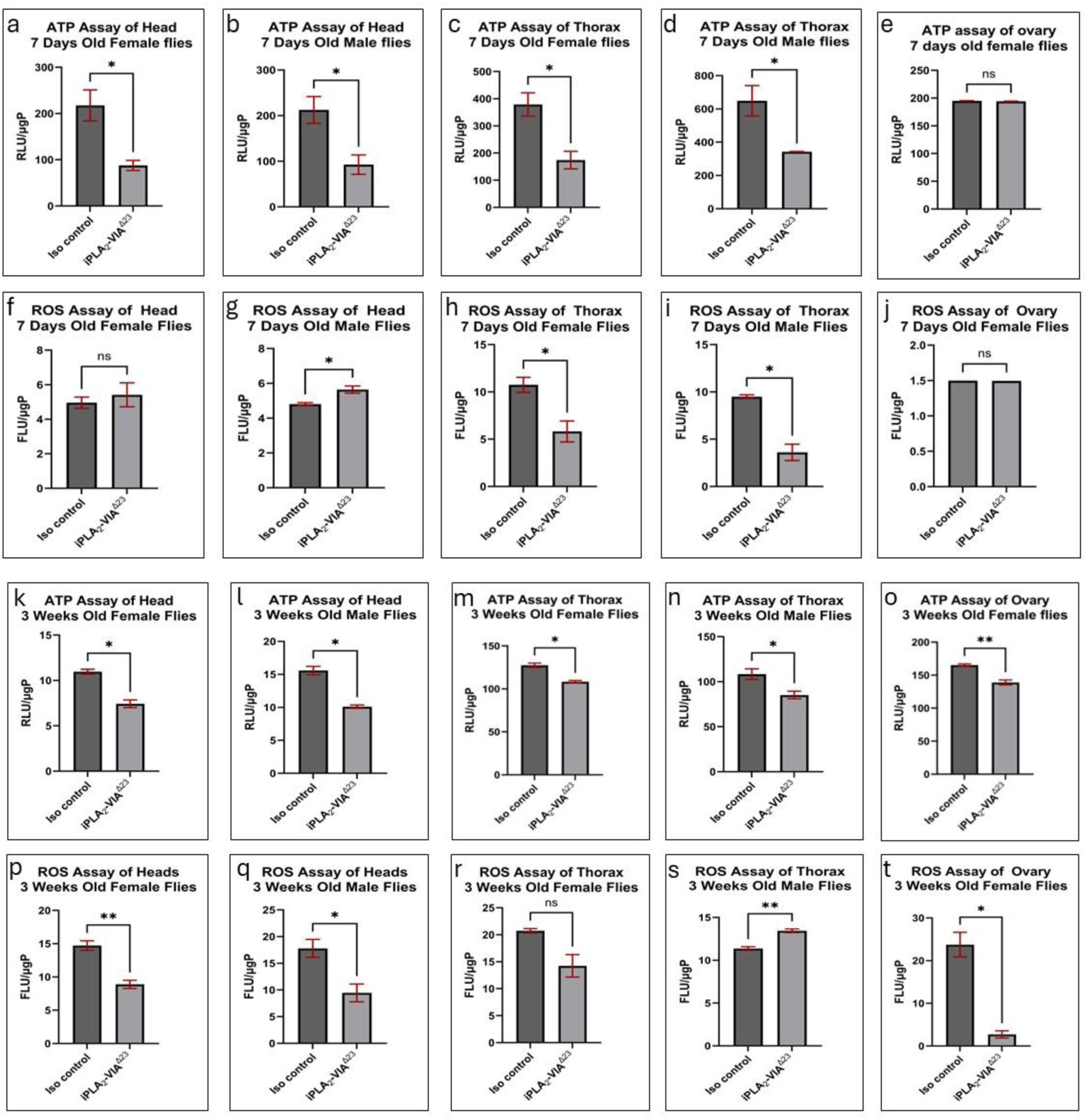
ATP production and reactive oxygen species (ROS) levels were measured in head, thorax, and ovary tissues from 7-day-old and 3-week-old iso-control and iPLA_2_-VIA mutant flies. Panels a–e show ATP levels in 7-day-old flies: female head (a), male head (b), female thorax (c), male thorax (d), and female ovary (e). Panels f–j show ROS levels in 7-day-old flies: female head (f), male head (g), female thorax (h), male thorax (i), and female ovary (j). Panels k–o show ATP levels in 3-week-old flies: female head (k), male head (l), female thorax (m), male thorax (n), and female ovary (o). Panels p–t show ROS levels in 3-week-old flies: female head (p), male head (q), female thorax (r), male thorax (s), and female ovary (t). ATP levels were significantly reduced in mutant heads and thoraxes at 7 days, while ovary ATP was not significantly changed at this age. At 3 weeks, ATP levels were significantly reduced in all examined mutant tissues. ROS levels showed tissue-, age-, and sex-dependent changes, with both decreases and increases observed in mutant tissues. ATP and ROS values were normalized to protein concentration and are presented as RLU/µg or FLU/µg protein, respectively. Data are shown as mean ± SEM. Statistical comparisons were performed between iso-control and iPLA_2_-VIA mutant groups within the same tissue, age, and sex using an unpaired two-tailed *t*-test.

In 7-day-old flies, ATP levels were significantly reduced in the heads of both female and male iPLA_2_-VIA mutants compared with iso-control flies (Figure 6a, b). This indicates that mitochondrial energy production is already impaired in young adult mutant neural tissue. ATP levels were also significantly reduced in the thoraxes of 7-day-old mutant females and males compared with their respective controls (Figure 6c, d), suggesting that muscle-associated mitochondrial function is also affected early in adulthood. In contrast, ATP levels in 7-day-old mutant ovaries were not significantly different from iso-control ovaries (Figure 6e), indicating that the functional effect on ATP production may be tissue-dependent at the young adult stage.

ROS levels in 7-day-old flies showed a more variable pattern than ATP. In 7-day-old female heads, ROS levels were not significantly different between iso-control and mutant flies (Figure 6f). However, in 7-day-old male heads, ROS levels were significantly increased in iPLA_2_-VIA *mutants* compared with controls (Figure 6g). In thoracic tissue, ROS levels were significantly reduced in both 7-day-old mutant females and males compared with controls (Figure 6h, i). In the ovary, ROS levels were not significantly changed in 7-day-old mutant flies (Figure 6j). These results show that ROS changes in young mutants are tissue- and sex-dependent, with increased ROS in male heads but reduced ROS in thoracic tissue.

In the 3-week-old flies, ATP production was significantly reduced across all examined mutant tissues. Aged iPLA_2_-VIA mutant females and males showed significantly lower ATP levels in the head compared with age-matched iso-controls (Figure 6k, l). ATP levels were also significantly reduced in the thoraxes of 3-week-old mutant females and males (Figure 6m, n). Unlike the young ovary, aged mutant ovaries showed a significant reduction in ATP production compared with iso-control ovaries (Figure 6o). These data indicate that ATP deficiency becomes more widespread with age, affecting neural, muscular, and reproductive tissues in aged mutant flies.

ROS levels in the 3-week-old flies also showed age-, sex-, and tissue-specific differences. In aged female and male heads, ROS levels were significantly reduced in iPLA_2_-VIA mutants compared with iso-control flies (Figure 6p, q). In the 3-week-old female thorax, ROS levels showed a decreasing trend but were not significantly different from controls (Figure 6r). In contrast, ROS levels were significantly increased in 3-week-old mutant male thorax compared with iso-control males (Figure 6s). In aged ovaries, ROS levels were strongly reduced in iPLA_2_-VIA mutants compared with controls (Figure 6t). Thus, ROS levels were not uniformly increased in mutant tissues; instead, they changed in a tissue-, sex-, and age-dependent manner.

Overall, ATP production was consistently reduced in iPLA_2_-VIA mutant flies, especially in head and thorax tissues at both ages and in the ovary at 3 weeks. This pattern suggests progressive impairment of mitochondrial bioenergetic capacity following loss of iPLA_2_-VIA. ROS levels showed a more complex pattern, with significant reductions in several tissues, no change in others, and increases in specific groups such as young male heads and aged male thorax. Together, these findings indicate that mitochondrial dysfunction in iPLA_2_-VIA mutants is characterized by reduced ATP production and disrupted redox homeostasis rather than a simple uniform increase in oxidative stress.

To further determine whether these tissue-specific alterations in mitochondrial function are reflected at the organismal level, we next quantified ATP production and ROS levels in whole adult flies. Consistent with the findings from individual tissues, iPLA_2_-VIA mutant flies exhibited a significant reduction in ATP levels across both ages and sexes, indicating a systemic impairment in mitochondrial energy production. ROS levels showed age- and sex-dependent changes, with reductions observed in most groups. These results demonstrate that mitochondrial dysfunction in iPLA_2_-VIA mutants is not restricted to specific tissues but represents a widespread defect affecting overall cellular energy homeostasis (Figure 7).

**Figure 7:**
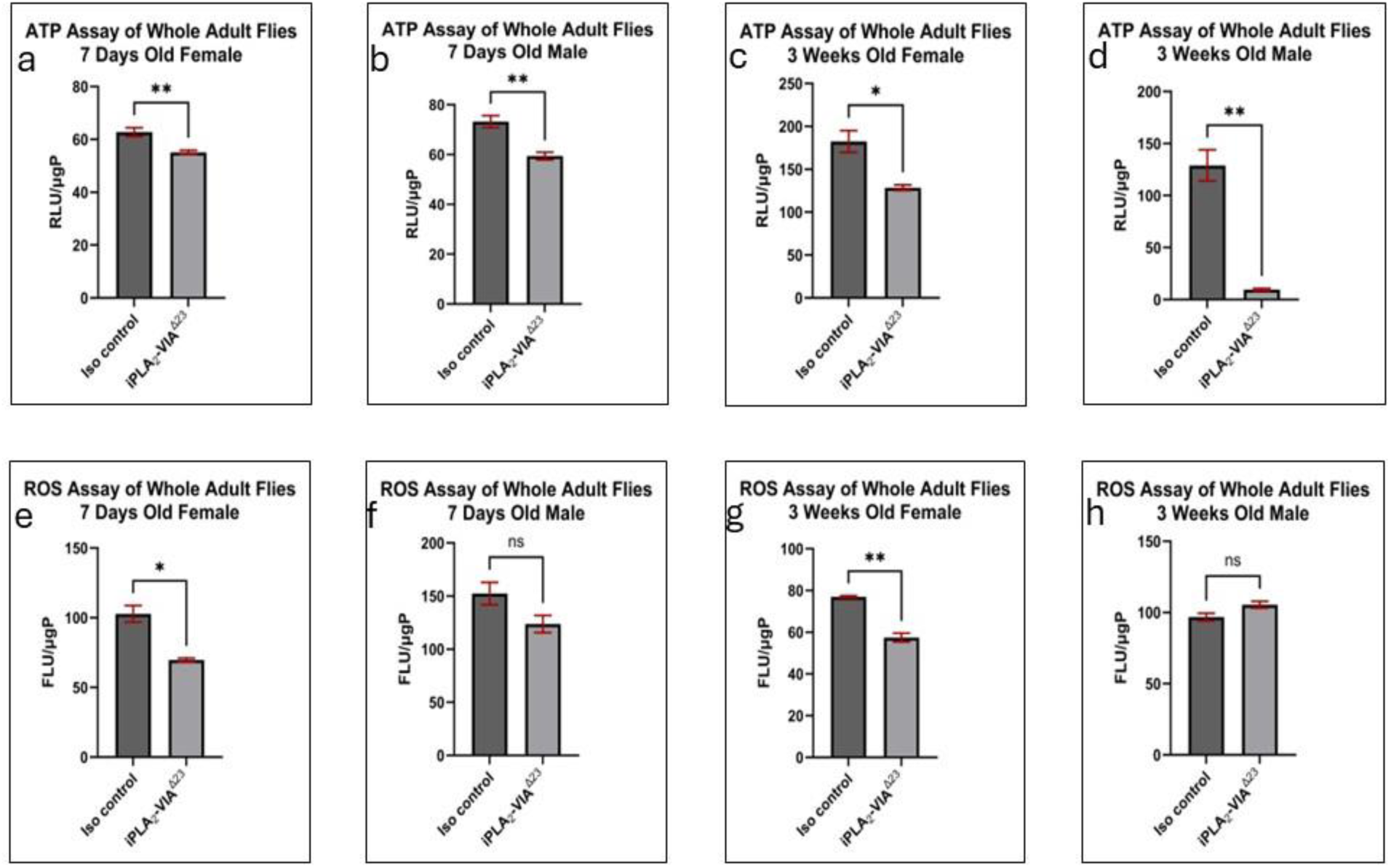
ATP production and reactive oxygen species (ROS) levels were measured in whole adult iso-control and iPLA_2_-VIA mutant *Drosophila melanogaster* at 7 days and 3 weeks of age. Panels a–d show ATP levels in 7-day-old females (a), 7-day-old males (b), 3-week-old females (c), and 3-week-old males (d). Panels e–h show ROS levels in 7-day-old females (e), 7-day-old males (f), 3-week-old females (g), and 3-week-old males (h). ATP levels were significantly reduced in mutant flies across both ages and sexes. ROS levels were significantly reduced in 7-day-old mutant females and 3-week-old mutant females, while no significant difference was observed in 7-day-old males or 3-week-old males. ATP and ROS values were normalized to protein concentration and are presented as RLU/µg or FLU/µg protein, respectively. Data are shown as mean ± SEM. Statistical comparisons were performed between iso-control and iPLA_2_-VIA mutant groups within the same age and sex using an unpaired two-tailed *t*-test.

Collectively, these results demonstrate that loss of iPLA_2_-VIA leads to progressive mitochondrial functional impairment, characterized by reduced ATP production and dysregulated ROS levels in an age- and sex-dependent manner. These functional deficits closely parallel the structural abnormalities and reduced mitochondrial number observed in mutant flies, providing a functional link between mitochondrial integrity and cellular energy homeostasis. This prompted us to examine next whether transcriptional regulators of mitochondrial biogenesis and dynamics are altered in iPLA_2_-VIA mutants.

### 6. Loss of iPLA2-VIA Selectively Reduces Mitochondrial-Encoded ATPase6 Transcript Expression

To determine whether loss of iPLA_2_-VIA affects the expression of ATP synthase genes encoded by mitochondrial DNA versus nuclear DNA, we measured transcript levels of *ATPase6* and *ATPSynC* by RT-qPCR. *ATPase6* is encoded by the mitochondrial genome and represents a mitochondrially encoded component of ATP synthase, whereas *ATPSynC* is encoded by the nuclear genome and represents a nuclear-encoded ATP synthase subunit. This comparison allowed us to determine whether ATP synthase-related gene expression was broadly altered or whether the effect was more specific to the mitochondrial genome.

In 7-day-old flies, *ATPase6* transcript levels were significantly reduced in iPLA_2_-VIA *mutants* of both sexes. Young mutant females showed a significant decrease in *ATPase6* expression compared with iso-control females (Figure 8a). Similarly, young mutant males also showed significantly reduced *ATPase6* expression compared with iso-control males (Figure 8b). These results indicate that mitochondrial-encoded ATP synthase gene expression is already reduced in young adult mutant flies.

**Figure 8:**
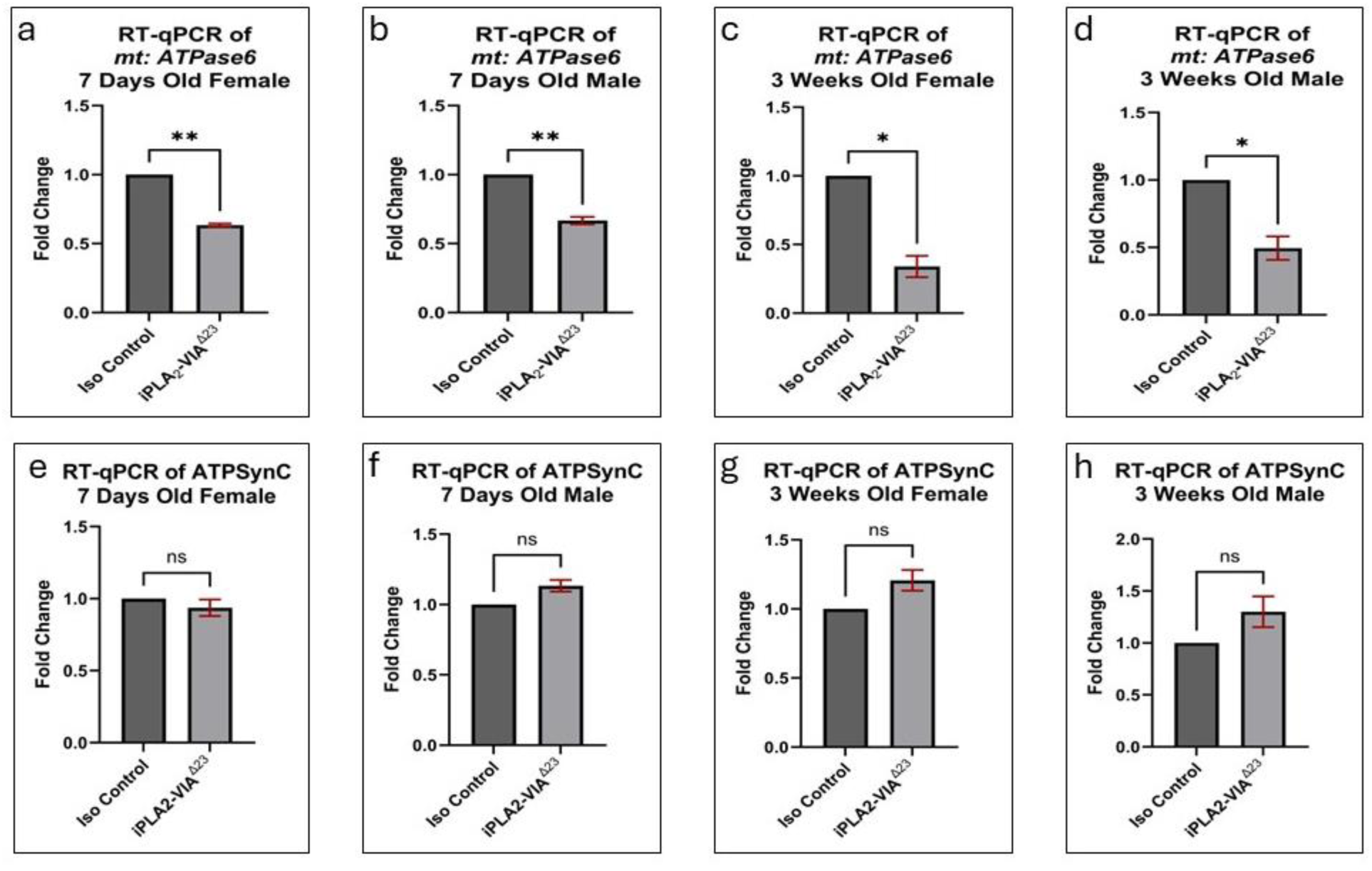
RT-qPCR analysis of *ATPase6* and *ATPSynC* transcript expression in iso-control and iPLA_2_-VIA mutant *Drosophila melanogaster* at 7 days and 3 weeks of age. Panels a–d show expression of the mitochondrial DNA-encoded ATP synthase gene *ATPase6* in 7-day-old females (a), 7-day-old males (b), 3-week-old females (c), and 3-week-old males (d). Panels e–h show expression of the nuclear DNA-encoded ATP synthase gene *ATPSynC* in 7-day-old females (e), 7-day-old males (f), 3-week-old females (g), and 3-week-old males (h). *ATPase6* transcript levels were significantly reduced in iPLA_2_-VIA mutant flies across both ages and sexes, whereas *ATPSynC* transcript levels were not significantly changed. Gene expression was normalized to *RpL32* and is presented as fold change relative to iso-control. Data are shown as mean ± SEM. Statistical comparisons were performed between iso-control and iPLA_2_-VIA mutant groups within the same age and sex using an unpaired two-tailed *t*-test.

Reduced *ATPase6* expression was also observed in aged flies. In 3-week-old females, iPLA_2_-VIA *mutants* showed a strong reduction in *ATPase6* transcript levels compared with age-matched iso-controls (Figure 8c). A significant decrease was also observed in 3-week-old mutant males compared with iso-control males (Figure 8d). Thus, reduced mitochondrial-encoded *ATPase6* expression was present across both ages and both sexes.

In contrast, the nuclear-encoded ATP synthase gene *ATPSynC* was not significantly altered in mutant flies. In 7-day-old females and males, *ATPSynC* transcript levels were not significantly different between iso-control and iPLA_2_-VIA *mutants* (Figure 8e, f). Similarly, in 3-week-old females and males, *ATPSynC* expression remained unchanged between control and mutant groups (Figure 8g, h).

Together, these results show that iPLA_2_-VIA mutant flies have reduced expression of the mitochondrial DNA-encoded ATP synthase gene *ATPase6*, while expression of the nuclear DNA-encoded ATP synthase gene *ATPSynC* remains largely preserved. This selective reduction in *ATPase6* transcript expression is consistent with the reduced mitochondrial DNA content observed in mutant flies and supports the conclusion that loss of iPLA_2_-VIA affects mitochondrial gene content and mitochondrial energy-related gene expression.

### 7. *Sod2* Expression Reveals Sex-Specific Alteration of Mitochondrial Antioxidant Defense

Since ROS levels were altered in an age-, sex-, and tissue-dependent manner in iPLA_2_-VIA mutant flies, we next examined whether mitochondrial antioxidant gene expression was also affected. We measured transcript levels of *Sod2*, which encodes mitochondrial superoxide dismutase, an antioxidant enzyme that helps regulate mitochondrial superoxide levels.

In 7-day-old female flies, *Sod2* expression was significantly reduced in iPLA_2_-VIA *mutants* compared with iso-control females (Figure 9a). This indicates that mitochondrial antioxidant defense is transcriptionally altered in young mutant females. In contrast, 7-day-old mutant males did not show a significant reduction in *Sod2* expression compared with iso-control males (Figure 9b).

**Figure 9:**
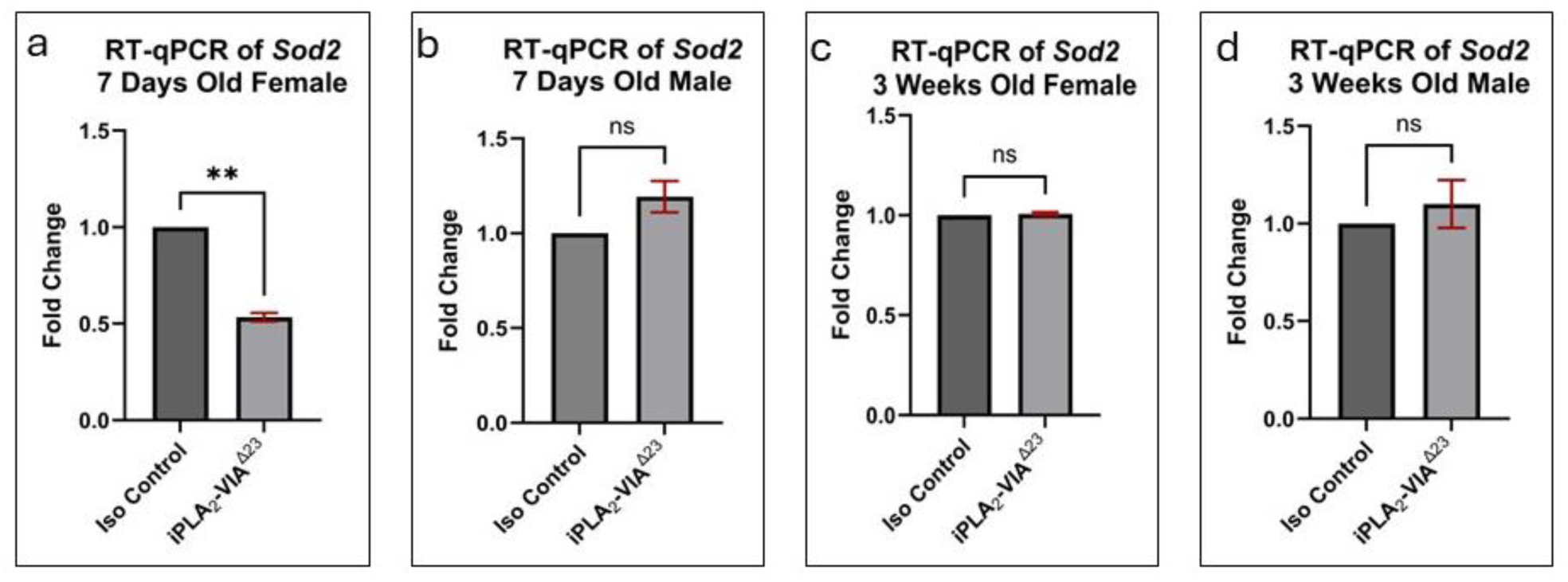
RT-qPCR analysis of *Sod2* transcript levels in iso-control and iPLA_2_-VIA mutant *Drosophila melanogaster* at 7 days and 3 weeks of age. Panels a–d show *Sod2* expression in 7-day-old females (a), 7-day-old males (b), 3-week-old females (c), and 3-week-old males (d). *Sod2* expression was significantly reduced in 7-day-old mutant females, while no significant difference was observed in 7-day-old males, 3-week-old females, or 3-week-old males. Gene expression was normalized to *RpL32* and is presented as fold change relative to iso-control. Data are shown as mean ± SEM. Statistical comparisons were performed between iso-control and iPLA_2_-VIA mutant groups within the same age and sex using an unpaired two-tailed *t*-test.

We then examined *Sod2* expression in aged flies. In 3-week-old females, *Sod2* transcript levels were not significantly different between iso-control and iPLA_2_-VIA mutant flies (Figure 9c). Similarly, 3-week-old mutant males showed no significant change in *Sod2* expression compared with iso-control males (Figure 9d).

Together with the ATP and ROS data, the selective reduction of *Sod2* in young mutant females supports the idea that iPLA_2_-VIA loss disrupts mitochondrial redox regulation rather than producing a uniform oxidative stress response across all groups.

Overall, *Sod2* expression was significantly reduced only in young female iPLA_2_-VIA *mutants*, while remaining largely unchanged in males and aged flies. This result suggests that changes in ROS levels in iPLA_2_-VIA mutants are not explained by a broad or consistent decrease in *Sod2* expression across all groups. Instead, *Sod2* changes appear to be sex- and age-specific, with the strongest effect observed in young mutant females.

### 8. iPLA_2_-VIA Loss Does Not Significantly Alter *Sirtuin 6* Expression

*Sirtuin 6* has been associated with mitochondrial regulation, metabolic homeostasis, and stress-response pathways. To determine whether iPLA_2_-VIA loss affects this regulatory pathway at the transcript level, we measured *Sirtuin 6* expression by RT-qPCR in 7-day-old and 3-week-old iso-control and iPLA_2_-VIA mutant flies of both sexes.

In 7-day-old females, *Sirtuin 6* expression was not significantly different between iso-control and iPLA_2_-VIA mutant flies (Figure 10a). Similarly, 7-day-old mutant males showed no significant change in *Sirtuin 6* transcript levels compared with iso-control males (Figure 10b). These results indicate that *Sirtuin 6* expression is not detectably altered in young adult mutant flies.

**Figure 10:**
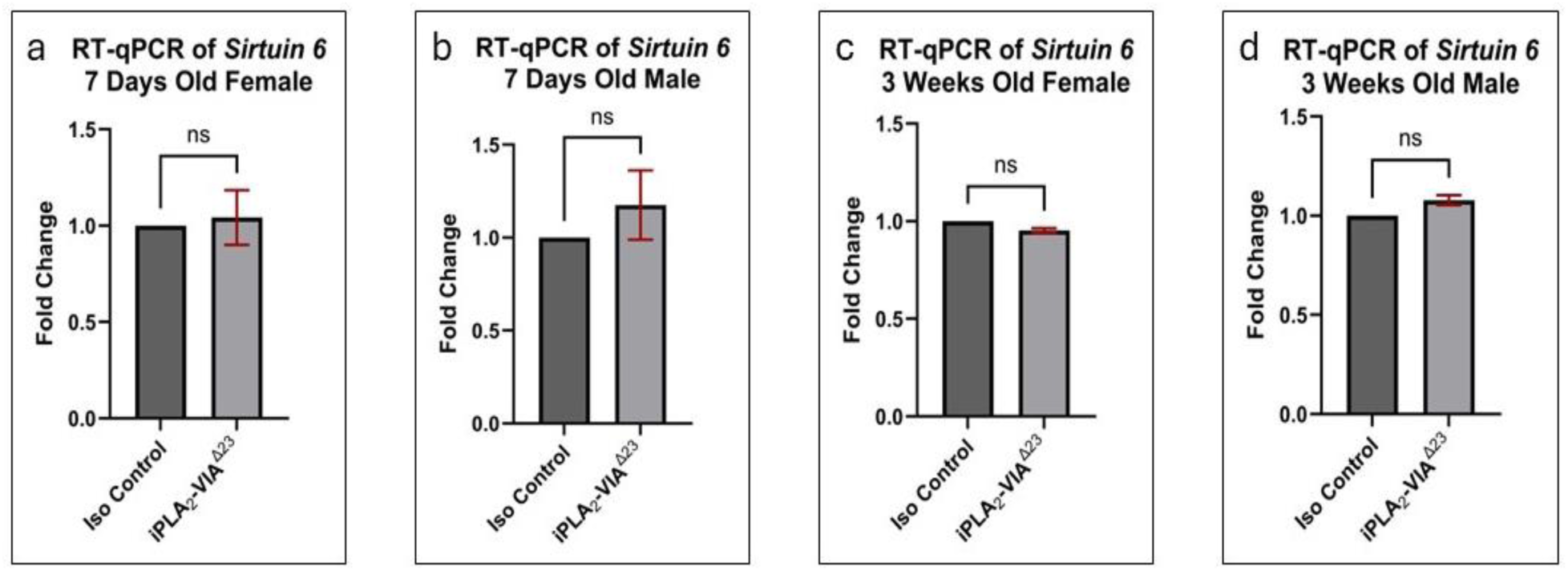
RT-qPCR analysis of *Sirtuin 6* transcript levels in iso-control and iPLA_2_-VIA mutant *Drosophila melanogaster* at 7 days and 3 weeks of age. Panels a–d show *Sirtuin 6* expression in 7-day-old females (a), 7-day-old males (b), 3-week-old females (c), and 3-week-old males (d). No significant differences in *Sirtuin 6* expression were observed between iso-control and iPLA_2_-VIA mutant flies in any age or sex group. Gene expression was normalized to *RpL32* and is presented as fold change relative to iso-control. Data are shown as mean ± SEM. Statistical comparisons were performed between iso-control and iPLA_2_-VIA mutant groups within the same age and sex using an unpaired two-tailed *t*-test.

We next examined aged flies. In 3-week-old females, *Sirtuin 6* expression remained unchanged between iso-control and iPLA_2_-VIA mutant flies (Figure 10c). Likewise, 3-week-old mutant males showed no significant difference in *Sirtuin 6* expression compared with age-matched iso-control males (Figure 10d).

Overall, *Sirtuin 6* transcript levels were not significantly altered in iPLA_2_-VIA mutant flies at either age or in either sex. This suggests that the mitochondrial structural and functional defects observed in the mutants are not accompanied by a detectable change in *Sirtuin 6* mRNA expression under the conditions examined.

### 9. *iPLA_2_-VIA* Loss Alters the Expression of Genes Regulating Mitochondrial Biogenesis and Dynamics

To check whether the reduction in mitochondrial number observed in iPLA_2_-VIA mutant flies was associated with altered mitochondrial biogenesis-related gene expression, we measured transcript levels of *mTOR* and *PGC1α* by RT-qPCR. These genes were selected because they are involved in cellular growth, metabolic regulation, and mitochondrial biogenesis.

In 7-day-old flies, *mTOR* expression was significantly reduced in both female and male iPLA_2_-VIA *mutants* compared with iso-control flies. Young mutant females showed lower *mTOR* transcript levels than iso-control females (Figure 11a). Similarly, young mutant males also showed significantly reduced *mTOR* expression compared with iso-control males (Figure 11b). These results show that *mTOR* expression is reduced early in adulthood in both sexes following loss of iPLA_2_-VIA.

**Figure 11:**
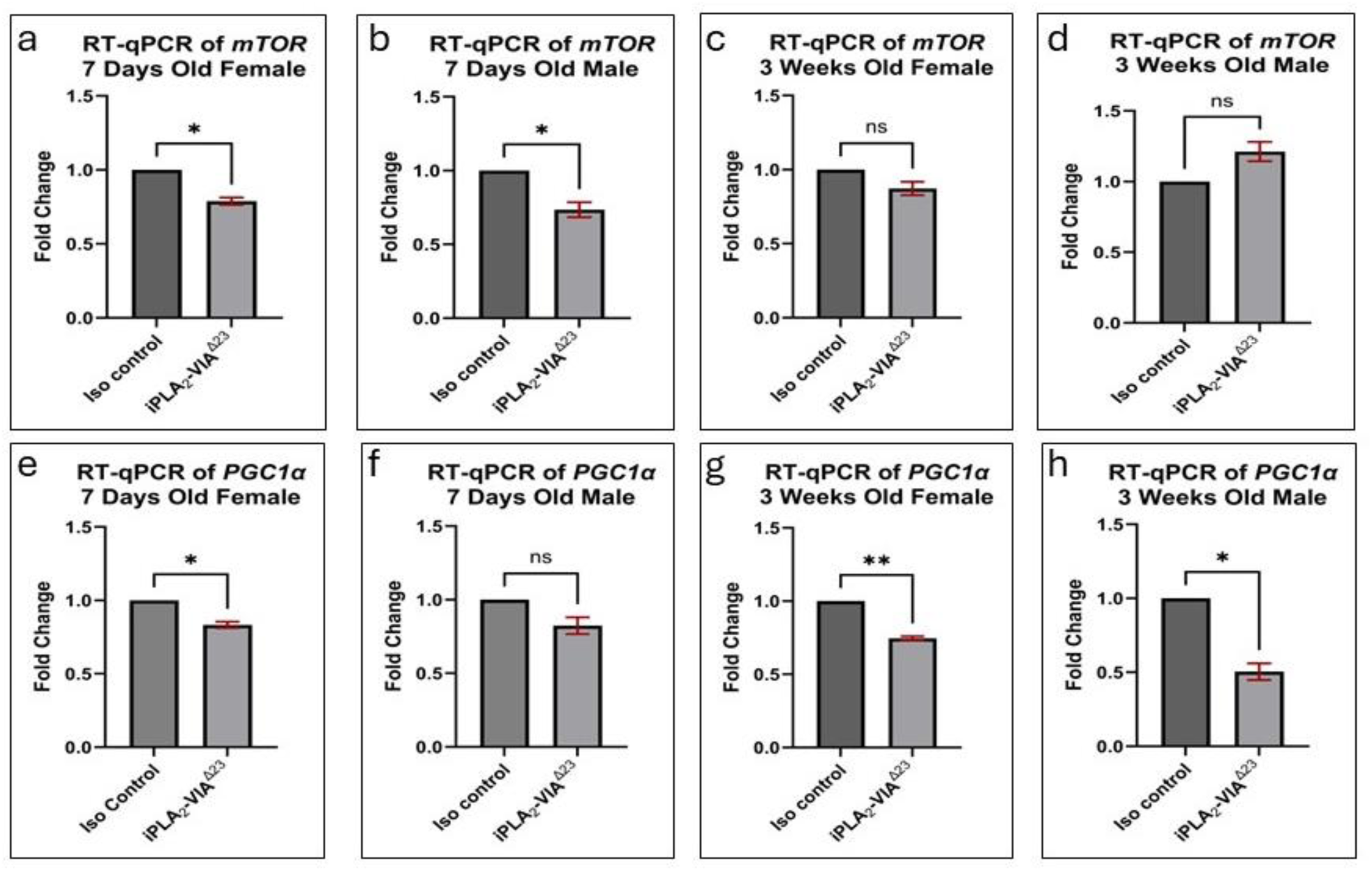
RT-qPCR analysis of *mTOR* and *PGC1α* transcript levels in iso-control and iPLA_2_-VIA mutant *Drosophila melanogaster* at 7 days and 3 weeks of age. Panels a–d show *mTOR* expression in 7-day-old females (a), 7-day-old males (b), 3-week-old females (c), and 3-week-old males (d). Panels e–h show *PGC1α* expression in 7-day-old females (e), 7-day-old males (f), 3-week-old females (g), and 3-week-old males (h). *mTOR* expression was significantly reduced in 7-day-old mutant females and males but was not significantly altered in 3-week-old mutants. *PGC1α* expression was significantly reduced in 7-day-old mutant females and in 3-week-old mutant females and males, while no significant change was observed in 7-day-old mutant males. Gene expression was normalized to *RpL32* and is presented as fold change relative to iso-control. Data are shown as mean ± SEM. Statistical comparisons were performed between iso-control and iPLA_2_-VIA mutant groups within the same age and sex using an unpaired two-tailed *t*-test.

In aged flies, the pattern of *mTOR* expression was different. In 3-week-old females, *mTOR* transcript levels were not significantly different between iso-control and iPLA_2_-VIA mutant flies (Figure 11c). Similarly, 3-week-old mutant males did not show a significant change in *mTOR* expression compared with age-matched iso-control males (Figure 11d). Therefore, the significant reduction in *mTOR* expression was mainly observed in young mutant flies, but not in aged mutants.

We next examined *PGC1α*, a key gene associated with mitochondrial biogenesis. In 7-day-old females, *PGC1α* expression was significantly reduced in iPLA_2_-VIA *mutants* compared with iso-control females (Figure 11e). In 7-day-old males, *PGC1α* expression showed a decreasing trend in mutants, but the difference was not statistically significant (Figure 11f). These data suggest that *PGC1α* expression is affected early in young mutant females, while the effect is less pronounced in young males.

In contrast to *mTOR*, *PGC1α* expression remained significantly altered in aged flies. In 3-week-old females, *PGC1α* transcript levels were significantly reduced in iPLA_2_-VIA *mutants* compared with iso-control females (Figure 11g). A similar significant reduction was observed in 3-week-old mutant males compared with iso-control males (Figure 11h). Thus, *PGC1α* expression was reduced in aged mutant flies of both sexes.

Following the analysis of mitochondrial biogenesis-associated genes, we next examined genes involved in mitochondrial fusion. Mitochondrial fusion is important for maintaining mitochondrial network organization, mitochondrial membrane integrity, and cristae structure. To determine whether loss of iPLA_2_-VIA affects fusion-related gene expression, we measured transcript levels of *Mfn1*, *Mfn2*, and *Opa1* in 7-day-old and 3-week-old iso-control and iPLA_2_-VIA mutant flies of both sexes (Figure 12).

**Figure 12:**
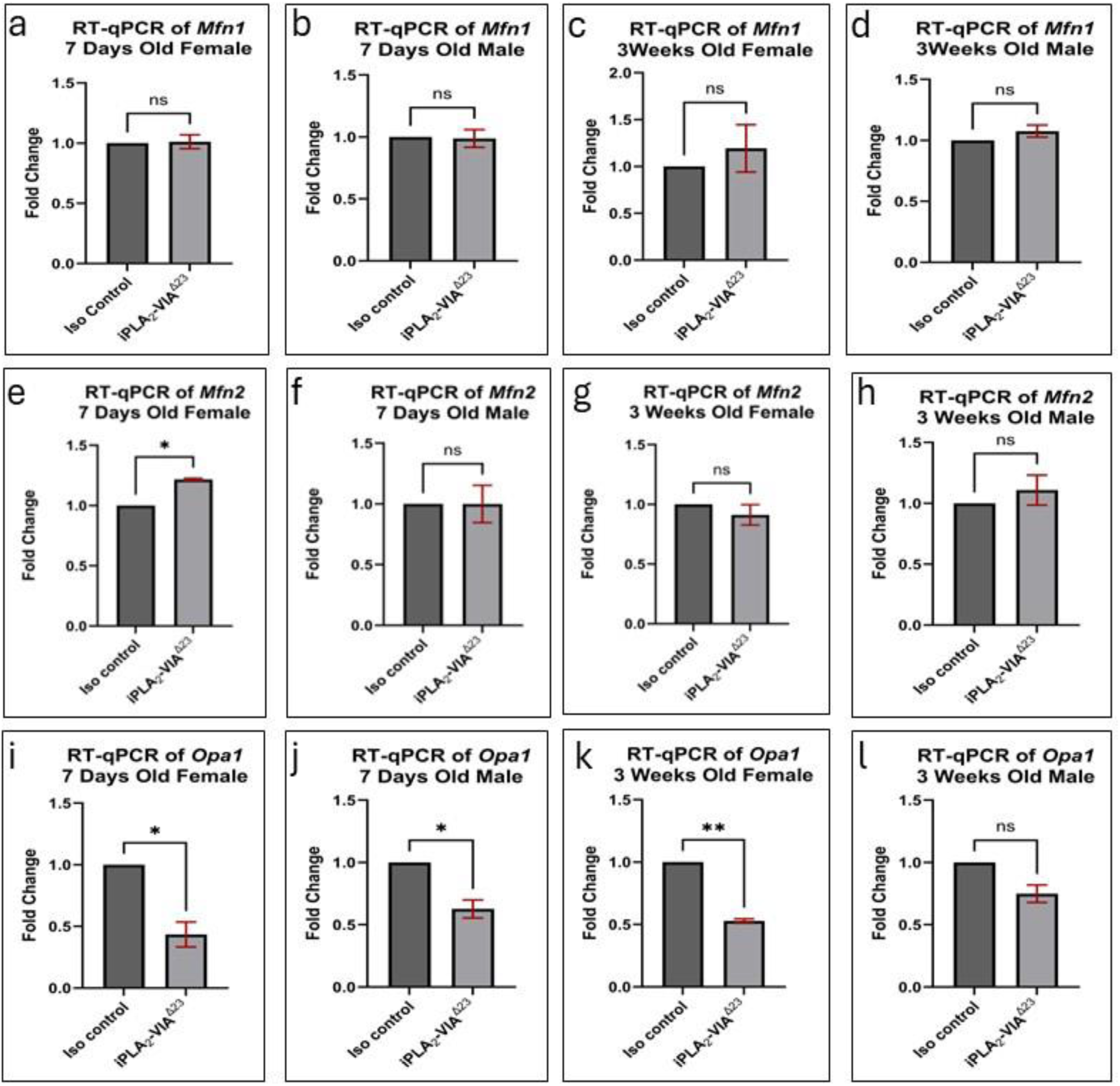
RT-qPCR analysis of mitochondrial fusion-associated genes *Mfn1*, *Mfn2*, and *Opa1* in iso-control and iPLA_2_-VIA mutant *Drosophila melanogaster* at 7 days and 3 weeks of age. Panels a–d show *Mfn1* expression in 7-day-old females (a), 7-day-old males (b), 3-week-old females (c), and 3-week-old males (d). Panels e–h show *Mfn2* expression in 7-day-old females (e), 7-day-old males (f), 3-week-old females (g), and 3-week-old males (h). Panels i–l show *Opa1* expression in 7-day-old females (i), 7-day-old males (j), 3-week-old females (k), and 3-week-old males (l). *Mfn1* expressions did not significantly change in any group. *Mfn2* expression was significantly increased only in 7-day-old mutant females, while no significant differences were observed in the other groups. *Opa1* expression was significantly reduced in 7-day-old mutant females, 7-day-old mutant males, and 3-week-old mutant females, but was not significantly changed in 3-week-old mutant males. Gene expression was normalized to *RpL32* and is presented as fold change relative to iso-control. Data are shown as mean ± SEM. Statistical comparisons were performed between iso-control and iPLA_2_-VIA mutant groups within the same age and sex using an unpaired two-tailed *t*-test.

*Mfn1* expression was not significantly altered in any age or sex group. In 7-day-old females, *Mfn1* transcript levels were similar between iso-control and iPLA_2_-VIA mutant flies (Figure 12a). No significant difference was observed in 7-day-old males either (Figure 12b). In aged flies, *Mfn1* expression also remained unchanged in both 3-week-old females and 3-week-old males (Figure 12c, d). These results indicate that iPLA_2_-VIA loss does not produce a detectable change in *Mfn1* transcript expression under the conditions examined.

We then measured *Mfn2* expression. In 7-day-old females, *Mfn2* transcript levels were significantly increased in iPLA_2_-VIA *mutants* compared with iso-control females (Figure 12e). However, this increase was not observed in 7-day-old males, where *Mfn2* expression was not significantly different between mutant and control flies (Figure 12f). In aged flies, *Mfn2* expression remained unchanged in both 3-week-old mutant females and 3-week-old mutant males compared with their respective iso-controls (Figure 12g, h). Therefore, the change in *Mfn2* expression was limited to young mutant females.

In contrast to *Mfn1* and *Mfn2*, *Opa1* expression showed a stronger reduction in iPLA_2_-VIA mutant flies. In 7-day-old females, *Opa1* transcript levels were significantly reduced in mutants compared with iso-control females (Figure 12i). A significant reduction was also observed in 7-day-old mutant males compared with iso-control males (Figure 12j). In aged flies, *Opa1* expression remained significantly reduced in 3-week-old mutant females (Figure 12k). However, in 3-week-old males, *Opa1* expression was lower in mutants but did not reach statistical significance (Figure 12l).

Following the analysis of mitochondrial fusion-associated genes, we examined whether loss of iPLA_2_-VIA also affects genes involved in mitochondrial fission. Mitochondrial fission is important for mitochondrial remodeling, distribution, and quality control. To assess fission-related transcriptional changes, we measured *Drp1* and *Fis1* expression by RT-qPCR in 7-day-old and 3-week-old iso-control and iPLA_2_-VIA mutant flies of both sexes.

In 7-day-old females, *Drp1* expression was strongly reduced in iPLA_2_-VIA *mutants* compared with iso-control females (Figure 13a). In contrast, 7-day-old mutant males showed a significant increase in *Drp1* expression compared with iso-control males (Figure 13b). This opposite pattern between young females and young males indicates that *Drp1* expression is altered in a sex-dependent manner during early adulthood.

**Figure 13:**
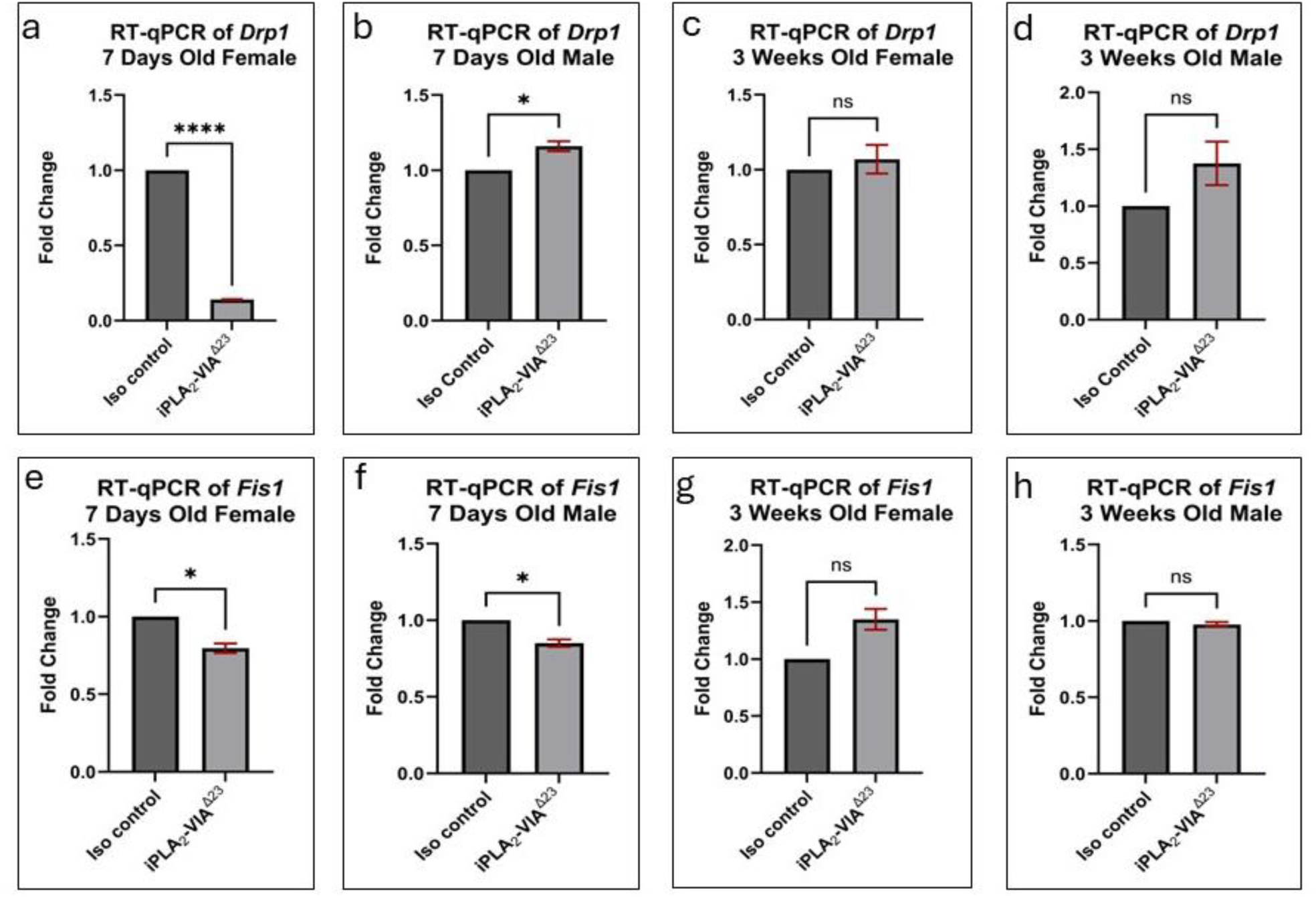
RT-qPCR analysis of mitochondrial fission-associated genes *Drp1* and *Fis1* in iso-control and iPLA_2_-VIA mutant *Drosophila melanogaster* at 7 days and 3 weeks of age. Panels a–d show *Drp1* expression in 7-day-old females (a), 7-day-old males (b), 3-week-old females (c), and 3-week-old males (d). Panels e–h show *Fis1* expression in 7-day-old females (e), 7-day-old males (f), 3-week-old females (g), and 3-week-old males (h). *Drp1* expression was significantly reduced in 7-day-old mutant females but significantly increased in 7-day-old mutant males, while no significant difference was observed in aged females or males. *Fis1* expression was significantly reduced in 7-day-old mutant females and males but was not significantly altered in 3-week-old mutants. Gene expression was normalized to *RpL32* and is presented as fold change relative to iso-control. Data are shown as mean ± SEM. Statistical comparisons were performed between iso-control and iPLA_2_-VIA mutant groups within the same age and sex using an unpaired two-tailed *t*-test

In aged flies, *Drp1* expression was not significantly altered. In 3-week-old females, *Drp1* transcript levels were similar between iso-control and iPLA_2_-VIA mutant flies (Figure 13c). Likewise, 3-week-old mutant males showed no significant difference in *Drp1* expression compared with age-matched iso-control males, although the mean expression appeared slightly higher in mutants (Figure 13d). These results suggest that significant *Drp1* dysregulation is mainly observed in young mutant flies.

We next examined *Fis1* expression. In 7-day-old females, *Fis1* transcript levels were significantly reduced in iPLA_2_-VIA *mutants* compared with iso-control females (Figure 13e). A similar significant reduction was observed in 7-day-old mutant males compared with iso-control males (Figure 13f). Thus, unlike *Drp1*, which changed in opposite directions between young females and males, *Fis1* was reduced in young mutants of both sexes.

In 3-week-old flies, *Fis1* expression was not significantly different between control and mutant groups. In aged females, *Fis1* expression showed no significant change between iso-control and iPLA_2_-VIA *mutants* (Figure 13g). Similarly, 3-week-old mutant males showed no significant difference in *Fis1* expression compared with iso-control males (Figure 13h).

We next examined *Pink1* expression to determine whether loss of iPLA_2_-VIA affects genes associated with mitochondrial quality control. *Pink1* is involved in the recognition and removal of damaged mitochondria, making it an important marker to examine in the context of mitochondrial structural defects, reduced mitochondrial number, and altered mitochondrial function observed in iPLA_2_-VIA mutant flies.

In 7-day-old flies, *Pink1* expression was significantly increased in both female and male iPLA_2_-VIA *mutants* compared with iso-control flies. Young mutant females showed higher *Pink1* transcript levels than iso-control females (Figure 14a). Similarly, young mutant males also showed significantly increased *Pink1* expression compared with iso-control males (Figure 14b). These findings indicate that *Pink1* expression is elevated early in adulthood in mutant flies of both sexes.

**Figure 14:**
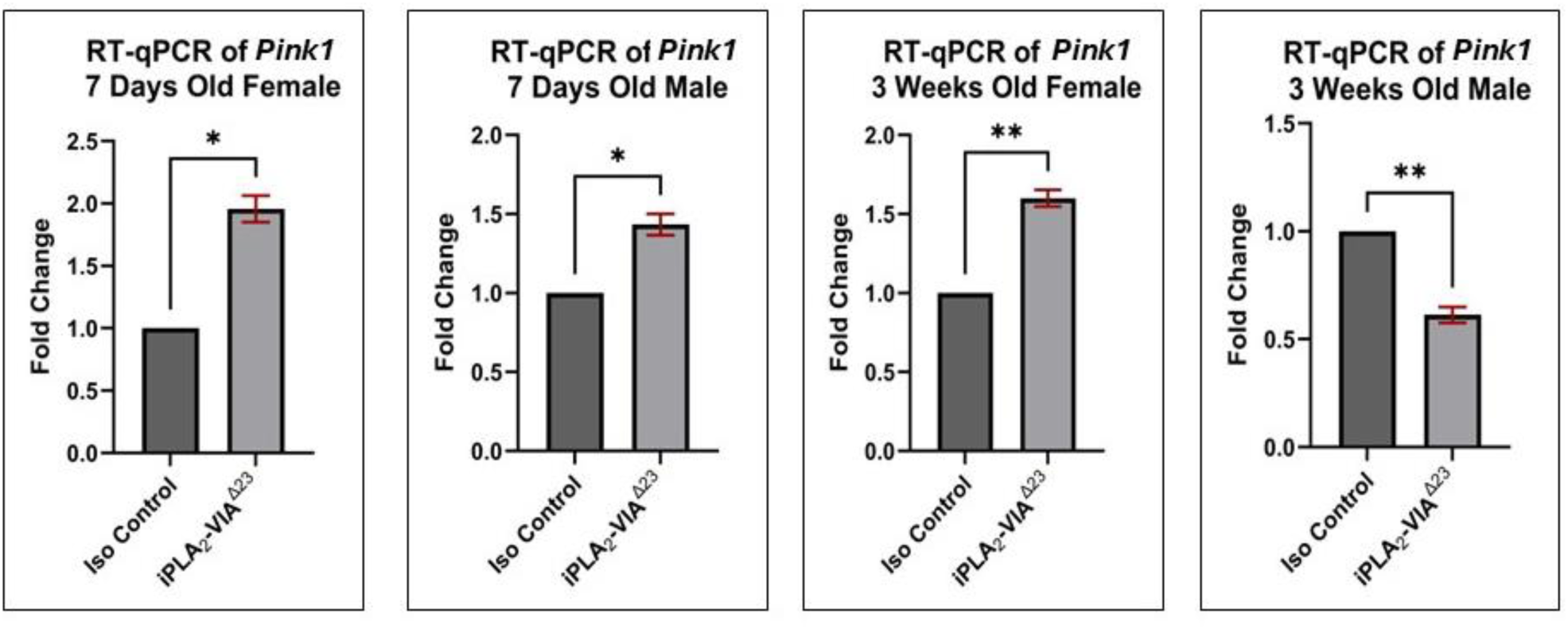
RT-qPCR analysis of *Pink1* transcript levels in iso-control and iPLA_2_-VIA mutant *Drosophila melanogaster* at 7 days and 3 weeks of age. Panels a–d show *Pink1* expression in 7-day-old females (a), 7-day-old males (b), 3-week-old females (c), and 3-week-old males (d). *Pink1* expression was significantly increased in 7-day-old mutant females, 7-day-old mutant males, and 3-week-old mutant females, whereas *Pink1* expression was significantly reduced in 3-week-old mutant males. Gene expression was normalized to *RpL32* and is presented as fold change relative to iso-control. Data are shown as mean ± SEM. Statistical comparisons were performed between iso-control and iPLA_2_-VIA mutant groups within the same age and sex using an unpaired two-tailed *t*-test.

In aged flies, *Pink1* expression showed a sex-dependent pattern. In 3-week-old females, *Pink1* expression remained significantly increased in iPLA_2_-VIA mutants compared with age-matched iso-control females (Figure 14c). In contrast, 3-week-old mutant males showed significantly reduced *Pink1* expression compared with iso-control males (Figure 14d). This opposite pattern in aged females and males suggests that *Pink1* regulation differs by sex during aging in iPLA_2_-VIA mutants.

Taken together with the altered expression of biogenesis (mTOR, PGC-1α), fusion (Opa1, Mfn2), fission (Drp1 and Fis1), and mitophagy-associated (PINK1) genes, these findings indicate that loss of iPLA2-VIA disrupts the coordinated transcriptional control of mitochondrial biogenesis, dynamics, and quality control, consistent with the structural abnormalities, reduced mitochondrial number, and impaired mitochondrial function observed in mutant flies.

### 10. *Trap1* Expression Declines in Old iPLA_2_-VIA Mutants

To further examine whether mitochondrial stress-response genes are altered following loss of iPLA_2_-VIA, we measured *Trap1* transcript levels by RT-qPCR in 7-day-old and 3-week-old iso-control and iPLA_2_-VIA mutant flies of both sexes. *Trap1* encodes a mitochondrial chaperone associated with mitochondrial protein homeostasis and stress protection.

In young flies, *Trap1* expression was not significantly altered in iPLA_2_-VIA mutants. In 7-day-old females, *Trap1* transcript levels were slightly higher in mutants compared with iso-control females, but this difference was not statistically significant (Figure 15a). Similarly, 7-day-old mutant males showed no significant change in *Trap1* expression compared with iso-control males (Figure 15b). These results indicate that *Trap1* expression is largely preserved in young mutant flies.

**Figure 15:**
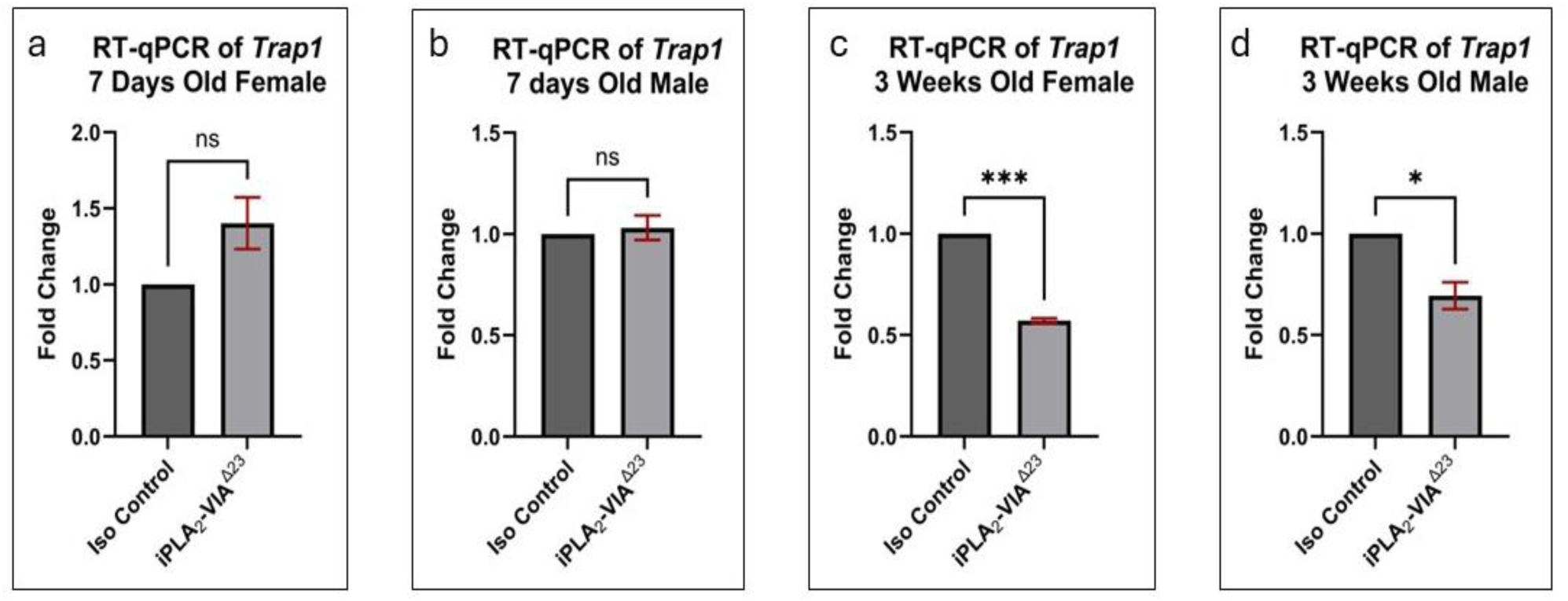
RT-qPCR analysis of *Trap1* transcript levels in iso-control and iPLA_2_-VIA mutant *Drosophila melanogaster* at 7 days and 3 weeks of age. Panels a–d show *Trap1* expression in 7-day-old females (a), 7-day-old males (b), 3-week-old females (c), and 3-week-old males (d). *Trap1* expression was not significantly changed in 7-day-old mutant females or males. In contrast, *Trap1* transcript levels were significantly reduced in 3-week-old mutant females and males compared with age-matched iso-controls. Gene expression was normalized to *RpL32* and is presented as fold change relative to iso-control. Data are shown as mean ± SEM. Statistical comparisons were performed between iso-control and iPLA_2_-VIA mutant groups within the same age and sex using an unpaired two-tailed *t*-test.

In aged flies, however, *Trap1* expression was significantly reduced in iPLA_2_-VIA *mutants*. In 3-week-old females, mutant flies showed a strong reduction in *Trap1* transcript levels compared with age-matched iso-control females (Figure 15c). A significant reduction was also observed in 3-week-old mutant males compared with iso-control males (Figure 15d). Thus, *Trap1* expression was specifically decreased in aged mutant flies of both sexes.

Overall, these data show that Trap1 expression is not significantly changed in young iPLA_2_-VIA mutants but is significantly reduced in aged mutants. This age-dependent decrease suggests that mitochondrial stress-response or chaperone-related gene expression becomes compromised during aging following loss of iPLA_2_-VIA. The downregulation of *Trap1* expression in aged flies is consistent with the more severe mitochondrial structural defects, reduced mitochondrial number, decreased ATP production, and altered redox status observed in older mutant flies.

## Discussion

In this study, we demonstrate that loss of function of iPLA_2_-VIA, the *Drosophila* homolog of human PLA2G6, leads to progressive and widespread mitochondrial degeneration and functional defects in both male and female mutant flies across multiple tissues. Mutant flies exhibited disrupted mitochondrial ultrastructure, reduced mitochondrial number, and mtDNA content. These defects ultimately lead to impaired ATP production and altered ROS homeostasis in the mutants. Age- and sex-dependent alterations in the expression of genes that regulate mitochondrial biogenesis, fusion-fission dynamics, and mitophagy in young and old mutant flies are likely to promote these mitochondrial defects in the mutants. Together, these findings indicate that disruption of iPLA_2_-VIA-dependent membrane lipid remodeling is sufficient to drive coordinated failure of mitochondrial production, maintenance and bioenergetic capacity in vivo in the mutants.

### Loss of iPLA_2_-VIA Disrupts Mitochondrial Ultrastructure Across Tissues

Previous studies have shown that PLA2G6 deficiency is associated with abnormal mitochondrial structure in old *Drosophila*, reduced mitochondrial membrane potential in the brain and ovary of old mutant flies, and impaired ATP production in the old mutant brain (Banerjee et al. 2021b; Kinghorn et al. 2015). However, whether these mitochondrial abnormalities occur across multiple tissues in an age- and sex-dependent manner has not been systematically examined. Our study extends these observations by showing that mitochondrial structural damage is not restricted to the brain but is also present in the thorax and ovary of young and old mutant flies (Figure 1a–t).

A central finding of this study is that iPLA_2_-VIA mutants exhibit severe mitochondrial ultrastructural abnormalities across the head, thorax, and ovary. In both 7-day-old and 3-week-old male and female mutant flies, mitochondria displayed reduced cristae density, disorganized cristae architecture, abnormal morphology, and compromised membrane integrity (Figure 1a–t). These structural abnormalities were accompanied by a significant reduction in mitochondrial number across multiple tissues, with reductions already present in several 7-day-old mutant tissues and becoming more widespread by 3 weeks of age (Figure 2a–j). These findings align with earlier reports of reduced mitochondrial membrane potential in iPLA_2_-VIA mutant flies (Banerjee et al. 2021b; Kinghorn et al. 2015).

These changes are functionally important because mitochondrial cristae provide the structural platform required for efficient oxidative phosphorylation and ATP production. Disruption of cristae organization can impair electron transport chain function and reduce ATP synthesis (Ježek et al. 2023). Thus, the defects observed by TEM are not merely morphological abnormalities; together with the reduction in mitochondrial number, they likely reflect loss of the mitochondrial membrane organization and abundance required for normal mitochondrial activity.

### Reduced Mitochondrial Number and mtDNA Content Indicate Mitochondrial Depletion

In addition to altered mitochondrial morphology, we observed a significant reduction in mitochondrial number in iPLA_2_-VIA mutant tissues, including the head/brain, thorax, and ovary. This decrease was evident in young mutant male and female flies and became more widespread with age, suggesting progressive mitochondrial depletion in the mutants (Figure 2a–j). Fluorescence imaging using MitoTracker Green further supported this finding in brain tissue. In 7-day-old mutant male and female brains, MitoTracker intensity appeared reduced compared with age-matched iso-control brains (Figure 3a–p). This reduction was also evident in 3-week-old mutant brains, where the mitochondrial signal appeared weaker compared to iso-control (Figure 4a–p). In particular, the 3-week-old mutant female brains showed a more punctate or clumpy MitoTracker staining pattern, suggesting abnormal mitochondrial distribution in addition to reduced mitochondrial signal (Figure 4e–h). This observation is relevant because a previous study in aged iPLA_2_-VIA mutant female germ cells reported abnormal mitochondrial aggregation, reduced mitochondrial membrane potential, and apoptosis (Banerjee et al. 2021b).

To determine whether reduced mitochondrial abundance was also reflected at the molecular level, we examined the relative levels of *ATPase6*, a mitochondrial DNA-encoded gene, and *ATPSynC*, a nuclear DNA-encoded gene, using DNA isolated from whole adult control and mutant flies of both sexes at young and aged stages. Consistent with the TEM quantification and MitoTracker imaging, mitochondrial DNA content was significantly decreased in young and aged mutant flies of both sexes, whereas nuclear DNA content remained unchanged (Figure 5a–h). These findings provide molecular evidence supporting reduced mitochondrial abundance in iPLA_2_-VIA mutants.

We also examined whether this reduction in mitochondrial DNA content was reflected at the transcript level. *ATPase6* transcript levels were significantly reduced in the whole body of young and aged male and female iPLA_2_-VIA mutants, while *ATPSynC* transcript levels were not significantly changed (Figure 8a–h). This selective reduction in mitochondrial-encoded *ATPase6* expression is consistent with the reduced mtDNA content observed in mutants and may indicate decreased mitochondrial gene output. However, because *ATPase6* transcript reduction could result from reduced mtDNA abundance, altered mitochondrial transcription, or both, this finding should be interpreted as evidence of reduced mitochondrial gene expression rather than direct proof of impaired mitochondrial transcription.

Together, these results indicate that loss of iPLA_2_-VIA leads to a selective reduction in mitochondrial abundance and mitochondrial genetic content rather than a broad loss of nuclear DNA. The combined evidence from TEM-based mitochondrial counting, reduced MitoTracker fluorescence, punctate or clumpy mitochondrial staining in aged mutant brains, decreased mtDNA content, and reduced *ATPase6* transcript expression supports the conclusion that iPLA_2_-VIA mutants undergo mitochondrial depletion. Since several mtDNA-encoded genes produce essential protein subunits of oxidative phosphorylation complexes, reduced mtDNA content may limit respiratory chain capacity and contribute to ATP deficits (Wang et al. 2021). This is particularly relevant for neuronal tissues, where mitochondrial ATP supports synaptic vesicle mobilization during exocytosis and recycling, axonal transport, calcium homeostasis, and long-term cell survival (Sheng and Cai 2012). Therefore, the combined reduction in mitochondrial number, MitoTracker signal, mtDNA content, and *ATPase6* expression in young and aged iPLA_2_-VIA mutant flies provides a molecular and cellular basis for impaired mitochondrial bioenergetics and increased vulnerability to neurodegeneration.

### Reduced ATP Production Reflects Impaired Mitochondrial Bioenergetic Capacity

The functional consequence of mitochondrial loss and structural damage was evident in ATP measurements. ATP levels were significantly reduced in the heads and thoraxes of 7-day-old male and female iPLA_2_-VIA mutant flies, while ATP levels in the ovary were not significantly altered at this young stage (Figure 6a–e). By 3 weeks of age, ATP levels were significantly reduced in all examined mutant tissues, including head, thorax, and ovary (Figure 6k–o). Whole-adult ATP measurements showed a similar pattern, with significantly reduced ATP levels in young and aged mutant males and females compared with iso-controls (Figure 7a–d). These findings indicate that loss of iPLA_2_-VIA causes broad mitochondrial bioenergetic impairment, which is already detectable in young flies and becomes more widespread with age.

Previous studies have shown that iPLA_2_-VIA mutants exhibit a severe age-dependent decline in locomotor ability (Banerjee et al. 2021b). Because neuronal and muscular tissues require high levels of ATP to sustain synaptic activity, ion homeostasis, muscle contraction, and tissue maintenance, reduced ATP production may contribute directly to locomotor impairment and neurodegenerative phenotypes in this model (Banerjee et al. 2021b; Sheng and Cai 2012). To our knowledge, however, ATP levels have not been systematically assessed across multiple tissues, ages, and sexes in iPLA_2_-VIA mutants. Our data addresses this gap by showing that bioenergetic impairment is widespread and progressively affects metabolically demanding tissues.

### ROS Changes Suggest Disrupted Redox Homeostasis Rather Than Uniform Oxidative Stress

ROS levels did not increase uniformly across tissues, ages, or sexes in iPLA_2_-VIA mutant flies. Instead, the mutants showed a complex pattern of ROS alteration. In 7-day-old flies, ROS levels were not significantly changed in female heads or ovaries, were significantly increased in male heads, and were significantly reduced in thoraxes of both sexes (Figure 6f–j). In 3-week-old flies, ROS levels were significantly reduced in female and male heads and in ovaries, showed no significant change in female thorax, and were significantly increased in male thorax (Figure 6p–t). Whole-adult ROS measurements also showed a sex-dependent pattern, with significant reductions in young and aged mutant females, but no significant change in young or aged mutant males (Figure 7e–h). Together, these results indicate that loss of iPLA_2_-VIA does not simply cause a generalized oxidative-stress response. Rather, it disrupts mitochondrial redox homeostasis in a tissue-, age-, and sex-dependent manner.

Mitochondrial defects, including cristae disruption, impaired membrane integrity, and reduced mtDNA content, can compromise electron transport chain activity (Li et al. 2025). In many cases, defective electron transport increases electron leakage and promotes ROS production (Jastroch et al. 2010). However, when mitochondrial damage is severe or respiratory activity is strongly reduced, ROS levels may also decrease because fewer electrons pass through the electron transport chain and fewer are available to react with oxygen (Addabbo et al. 2009). Therefore, reduced ROS in some mutant tissues may reflect diminished mitochondrial respiratory capacity rather than improved redox status. This interpretation is consistent with our ATP data, which showed reduced bioenergetic output in mutant tissues and whole flies (Figures 6a–e, 6k–o, and 7a–d). In contrast, tissues showing increased ROS, such as young mutant male heads and aged mutant male thorax, may represent conditions in which damaged mitochondria remain metabolically active but inefficient, leading to electron leakage and oxidative imbalance.

It is also important to note that ROS are not only damaging molecules. At low to moderate levels, ROS act as signaling molecules that regulate neuronal function, synaptic plasticity, stress responses, and cellular adaptation (Massaad and Klann 2011). Therefore, both excessive ROS and abnormally reduced ROS can disturb normal cellular signaling. In this context, the altered ROS patterns observed in iPLA_2_-VIA mutants likely reflect disruption of redox balance rather than a simple increase in oxidative stress.

### *Sod2* Expression Suggests Limited and Context-Dependent Antioxidant Gene Response

Because ROS levels were altered in a tissue-, age-, and sex-dependent manner, we examined *Sod2* expression to determine whether mitochondrial antioxidant defense was also affected. *Sod2* is the major mitochondrial matrix enzyme that converts superoxide radicals into hydrogen peroxide, thereby limiting oxidative damage within mitochondria (Islam et al. 2026). Previous studies have shown that loss or strong knockdown of *Sod2* can cause precocious neurodegeneration and DNA strand breaks in neurons (Paul et al. 2007).

In our study, *Sod2* expression was significantly reduced only in 7-day-old mutant females, whereas no significant change was observed in 7-day-old mutant males or in aged mutant flies of either sex (Figure 9a–d). This selective pattern suggests that ROS alterations in iPLA_2_-VIA mutants are not driven primarily by broad transcriptional suppression of *Sod2*. Instead, the altered ROS profile likely reflects a combination of reduced mitochondrial number, disrupted cristae structure, decreased mtDNA content, and impaired respiratory activity. In tissues where mitochondrial respiratory capacity is strongly reduced, ROS production may decrease because fewer electrons pass through the electron transport chain. In contrast, tissues with damaged but still metabolically active mitochondria may show increased ROS due to inefficient electron transfer.

Together with the ATP and ROS results, the *Sod2* data support the idea that loss of iPLA_2_-VIA disrupts mitochondrial redox regulation rather than causing a uniform oxidative stress response. The sex-specific decrease in *Sod2* also suggests that female mutants may engage different or insufficient antioxidant responses during early mitochondrial stress, which could contribute to the tissue- and sex-dependent mitochondrial phenotypes observed in this model.

### *Sirtuin 6* Expression Is Preserved Despite Mitochondrial Dysfunction

Although *SIRT6* has been reported to regulate mitochondrial function, particularly in metabolically active tissues such as the brain (Smirnov et al. 2023). However, in our study, *Sirtuin 6* transcript levels were not significantly changed in iPLA_2_-VIA mutant flies at either age or in either sex (Figure 10a–d). This suggests that the mitochondrial abnormalities observed in this study are unlikely to be driven by broad transcriptional alteration of *Sirtuin 6*. However, because *Sirt6* function can also be regulated at the protein, enzymatic, or post-translational level, unchanged mRNA expression does not exclude altered *Sirt6* activity. Future studies measuring *Sirt6* protein abundance or activity would be needed to determine whether this pathway contributes to mitochondrial dysfunction in iPLA_2_-VIA mutants.

Instead, the defects appear more closely associated with reduced mitochondrial content, impaired *mTOR–PGC-1α* biogenesis signaling, altered fusion–fission gene expression, and disrupted mitochondrial quality-control pathways.

### Impaired Mitochondrial Biogenesis and Enhanced Mitophagy of Damaged Mitochondria

Both mTOR and PGC-1α are important regulators of mitochondrial biogenesis, with mTOR acting upstream to coordinate nutrient and energy status with mitochondrial growth, and PGC-1α functioning as a central transcriptional co-activator that promotes mitochondrial gene expression, oxidative phosphorylation, and antioxidant defense (Rius-Pérez et al. 2020; Summer et al. 2019). In our study, *Mtor* expression was significantly reduced in young iPLA_2_-VIA mutant males and females, while *PGC-1α* expression was reduced in young mutant females and in aged mutant flies of both sexes (Figure 11a–h). These findings suggest that mitochondrial biogenesis-associated signaling is altered following loss of iPLA_2_-VIA. Previous studies have also shown that reduced mTOR function increases lifespan, reduces female fertility (Guo and Yu 2019; Papadopoli et al. 2019), impairs climbing ability in flies (Birse et al. 2010), promotes mitophagy and thereby reduces ROS production (Krishnamoorthy et al. 2018; Weichhart 2018). Accordingly, iPLA_2_-VIA mutant flies exhibited female specific fertility defects, age dependent climbing defect (Banerjee et al. 2021b), and reduced ROS production (Figure 7 e-h) suggesting that reduced *mTOR* expression in PLAN flies may trigger these physiological effects, whereas, the reduction of *mTOR* expression in them may be driven by a feedback mechanism to promote mitophagy and rescue their severely reduced lifespan (Banerjee et al. 2021b). Because PGC-1α is a major regulator of mitochondrial biogenesis and metabolic homeostasis, its downregulation is consistent with the reduced mitochondrial number and decreased mtDNA content observed in mutant flies. Suppression of the mTOR–PGC-1α axis would be expected to alter nutrient and energy homeostasis in the mutant flies, as this signaling pathway tightly regulates nutrient sensing, storage, mobilization and utilization for energy production (Goul et al. 2023).

### Altered Expression Levels of Fusion and Fission Genes Indicate Disrupted Mitochondrial Dynamics

We also observed selective changes in the expression of the genes regulating mitochondrial fusion and fission dynamics. Among the major mitochondrial fusion-related genes, *Mfn1* expression was largely preserved, while *Mfn2* showed limited sex-specific changes, and *Opa1* was significantly reduced in young and old male and female mutant flies (Figure 12a–l). Because Opa1 plays an important role in inner mitochondrial membrane fusion and cristae maintenance (Patten et al. 2014), the reduced *Opa1* expression explains the disrupted cristae architecture, reduced cristae density, and abnormal mitochondrial morphology observed by TEM in the mutants. This interpretation is supported by previous studies showing that heterozygous *Opa1* mutants have shortened lifespan and abnormal mitochondrial structure in muscle tissue (Tang et al. 2009), which overlap with our observations of mitochondrial structural defects, and previous studies that reported markedly reduced lifespan of iPLA_2_-VIA homozygous null mutant flies (Banerjee et al. 2021b).

In parallel, the fission-related genes *Drp1* and *Fis1* showed age- and sex-dependent dysregulation in the PLAN flies. Both genes were significantly altered in young mutant males and females, whereas these differences were no longer significant at the older age (Figure 13a–h). This pattern suggests that mitochondrial fission pathways are disrupted early after loss of iPLA_2_-VIA, but may become compensated, or masked by later-stages of mitochondrial loss and tissue degeneration. Drp1 is a central GTPase that mediates mitochondrial fission and plays an important role in mitochondrial quality control, mitophagy, calcium handling, and apoptosis (Cai et al. 2024). Proper Drp1 activity allows damaged mitochondrial regions to be separated and removed, whereas excessive or insufficient fission can disrupt mitochondrial network integrity (Nivedya et al. 2025). Fis1 is an outer mitochondrial membrane protein that helps regulate fission through its interaction with Drp1. Reduced Fis1 activity has been shown to lower mitochondrial membrane potential, suggesting an important role for Fis1 in maintaining mitochondrial membrane integrity and bioenergetic function (Strucinska et al. 2025). In our study, altered *Fis1* expression in iPLA_2_-VIA mutants coincides with reduced mitochondrial membrane potential (Banerjee et al. 2021b; Kinghorn et al. 2015), supporting the idea that disrupted fission-related signaling may contribute to mitochondrial functional decline.

### *Pink1* Dysregulation Suggests Altered Mitochondrial Quality Control

The altered expression of *Pink1* adds another layer to the mitochondrial phenotype. Pink1 is a key regulator of mitophagy and promotes the removal of damaged mitochondria (Quinn et al. 2020). Increased *Pink1* expression in young mutants and old mutant females may reflect activation of mitochondrial quality-control signaling in response to mitochondrial damage. However, reduced *Pink1* expression in old mutant males suggests that this response may be sex- and age-dependent and may fail in certain contexts (Figure 14). One possible interpretation is that early mitochondrial damage activates mitophagy-related signaling in the young mutants, but this response becomes insufficient or dysregulated as damage accumulates with age in the old mutant males only.

### Age-Dependent *Trap1* Reduction Suggests Impaired Mitochondrial Stress Protection

The age-dependent reduction of *Trap1* expression in iPLA_2_-VIA mutants is particularly relevant in light of previous work connecting Trap1 to mitochondrial protection in *Drosophila* models of Parkinson’s disease. A previous study showed that loss of Trap1 decreases mitochondrial function, reduces ATP levels, increases stress sensitivity, and lowers dopamine content in fly heads. They further demonstrated that neuronal upregulation of *Trap1* can rescue mitochondrial impairment in *Pink1* mutants, placing *Trap1* downstream of *Pink1* and in parallel with Parkin in maintaining mitochondrial function (Costa et al. 2013). In our study, *Trap1* expression was significantly reduced only in old iPLA_2_-VIA mutants (Figure 15a–d), coinciding with severe mitochondrial structural defects, reduced mitochondrial number, mtDNA depletion, and reduced ATP production. An earlier study in *Trap1* mutant flies showed that respiration rate by mitochondrial electron transport complexes I and II was severely reduced, like in iPLA_2_-VIA mutant flies (Kinghorn et al. 2015), further adding to the molecular basis of mitochondrial functional impairment in the iPLA_2_-VIA mutant old male and female flies. Moreover, dopaminergic neurons degenerate progressively in the PLA2G6 knock-out mice brain (Beck et al. 2016), and loss of Trap1 accelerates the dopaminergic neuron degeneration in flies (Costa et al. 2013), which is responsible for their disabled locomotion ability (Mori et al. 2019). Although we did not directly examine dopaminergic neurons in iPLA_2_-VIA mutants, our overall results imply that severe climbing defects in 3-week-old iPLA_2_-VIA mutants (Banerjee et al. 2021b) resulted from the loss of dopaminergic neurons in their brain due to downregulation of *Trap1*.

## Conclusion

In conclusion, this is the first study demonstrating that loss of iPLA_2_-VIA causes coordinated mitochondrial dysfunction across multiple tissues in *Drosophila melanogaster*. Mutant flies exhibit widespread mitochondrial ultrastructural abnormalities, reduced mitochondrial number, decreased mtDNA content, impaired ATP production, and altered ROS homeostasis. These defects appear early in adulthood and become more pronounced with age, indicating that mitochondrial impairment is a progressive consequence of iPLA_2_-VIA loss. Additionally, we show that the mitochondrial genome number is also reduced in the mutants. This mitochondrial depletion in the mutants is accompanied by the loss of its biogenesis due to downregulation of mTOR–PGC-1α signaling, and by the enhanced mitophagy due to downregulation of *mTOR* and upregulation of *Pink1* expression. The altered expression of fusion and fission regulatory genes indicates disruption of mitochondrial maintenance and quality-control in the iPLA_2_-VIA mutant flies. Decline in fertility and locomotor activity is triggered by the downregulation of *mTOR* and *Trap1* expression in the mutants. The old mutant flies are likely to lose dopaminergic neurons in their brain as a result of downregulation of *Trap1*.

This combined failure of mitochondrial structure, abundance, function, and quality control provides a mechanistic framework for understanding how *PLA2G6/iPLA_2_-VIA* deficiency contributes to progressive neurodegenerative phenotypes. These findings highlight mitochondrial maintenance pathways, including mitochondrial biogenesis, fusion–fission balance, redox regulation, and mitophagy, as important targets for future studies aimed at identifying therapeutic strategies for PLAN and related neurodegenerative disorders.

